# PD1-induced Shp2 condensation organizes inhibitory signalosomes through selective substrate partitioning

**DOI:** 10.64898/2026.03.09.710629

**Authors:** Takeya Masubuchi, George A Wen, Xiaoxian Song, Keshav Gaddam, Haian Shao, Chuan Wu, Enfu Hui

## Abstract

The formation of microclusters is a hallmark of PD1 engagement with its ligands, yet the physical basis and functional significance of this phenomenon remain unclear. Here we show that ligand-bound PD1 licenses Shp2 self-association and liquid-liquid phase separation (LLPS), producing dynamic PD1:Shp2 condensates whose liquidity depends on Shp2 catalytic activity. Mutations that selectively disrupt Shp2 self-association weaken PD1 microcluster formation and impair PD1 inhibitory function. Mechanistically, PD1-induced Shp2 LLPS promotes the co-compartmentalization of signaling substrates such as CD3ζ and CD28, thereby facilitating its dephosphorylation. These findings identify Shp2 LLPS as an intrinsic organizing principle of the PD1 inhibitory pathway that links enzymatic activation, substrate selectivity and mesoscale assembly to suppress T cell activation.

## INTRODUCTION

Programmed cell death protein 1 (PD1) is an immune checkpoint receptor expressed on multiple immune cell types, where it suppresses immune activation to maintain immune homeostasis (*1, 2*). PD1 is highly expressed on exhausted tumor-infiltrating T cells and limits their anti-tumor responses. Blockade therapy targeting PD1 or its ligand PDL1 has achieved remarkable clinical success in some patients (*3*). However, the overall response rates to PD1 blockade remain limited across many cancer types, underscoring the need for a deeper understanding of PD1 biology and signaling.

PD1 is a type I transmembrane protein consisting of an IgV-like extracellular domain, a single-pass transmembrane segment, and an intracellular domain (ICD) containing an immunoreceptor tyrosine-based inhibitory motif (ITIM) and an immunoreceptor tyrosine-based switch motif (ITSM) (*4, 5*). Upon binding to its ligands PD-L1 (*6, 7*) or PD-L2 (*8*), PD1 becomes phosphorylated at the tyrosine residues within ITIM and ITSM, leading to the recruitment of cytosolic protein tyrosine phosphatase (PTPase) Shp2 (*4, 5, 9*). PD1-associated Shp2 then dephosphorylates proximal signaling molecules, thereby attenuating T cell activation. During this process, PD1 forms microscale clusters, or microclusters, that enrich Shp2 and colocalize with T cell antigen receptors (TCR) and costimulatory receptors (*10–13*). While PD1 microcluster formation has been considered a hallmark of PD1 inhibitory signaling (*14*), the driving force and functional significance of these assemblies remain poorly understood.

Cellular signal transduction depends on the interplay between biochemical reactions and the spatial organization of signaling proteins. A transformative mechanism for this spatial organization is liquid–liquid phase separation (LLPS) (*15–20*), whereby multivalent interactions drive the formation of dynamic, membraneless condensates that locally concentrate signaling molecules (*15, 17*). LLPS has emerged as a general principle underlying the assembly of many signaling complexes, including the TCR–LAT signalosome (*17*), cGAS–STING clusters (*21*), and Ras nanoclusters (*22*). Whether a similar mechanism contributes to the organization of PD1 signaling remains unknown.

Shp2, the principal effector recruited to PD1, undergoes dynamic conformational changes upon binding to phosphotyrosine (pY) peptides. Shp2 consists of two Src homology 2 (SH2) domains, an N-terminal SH2 (nSH2) and a C-terminal SH2 (cSH2), followed by a single PTPase domain. In its basal state, Shp2 adopts a closed, autoinhibited conformation in which the nSH2 domain interacts with the PTPase domain, occluding the catalytic pocket from accessing its substrates (*23, 24*). Upon binding to pY peptides via its SH2 domains, Shp2 transitions into an open conformation, where the nSH2 domain relocates to the opposite surface of the PTPase domain, thereby exposing the active site and enabling the catalytic activity (*25–28*).

LLPS has been described as a feature of certain disease-associated mutants of Shp2 (*24*), which promote uncontrolled cell growth causing cancer and developmental syndromes (*29–32*). Namely, these Shp2 mutants spontaneously undergo LLPS via intermolecular electrostatic interactions between the basic and acidic patches on its PTPase domain (*24*). However, in wild-type (WT) Shp2 (denoted as Shp2^WT^ herein), the basic patches in the PTPase domain are normally shielded by the nSH2 in *cis*, thereby restricting the intermolecular (*trans*) associations of the PTPase domains and LLPS (*24*). Whether the LLPS potential of the Shp2 PTPase domain is integral to normal Shp2 function and whether it regulates PD1 function is still unknown.

Here, we show that PD1 engagement unleashes the latent LLPS potential of Shp2, promoting its self-association and condensation into PD1:Shp2 assemblies with liquid-like properties. These condensates act as biochemical microreactors that selectively enrich ICDs of CD3ζ and CD28 while excluding that of the inhibitory receptor TIGIT (*33, 34*), thereby organizing an inhibitory signalosome optimized for substrate access and dephosphorylation.

Using biochemical reconstitution, live-cell imaging, and T-cell functional assays, we demonstrate that disrupting Shp2 LLPS impairs PD1 microcluster formation, substrate co-localization, and inhibitory signaling. Our findings reveal that PD1-induced Shp2 condensation integrates enzymatic activation with substrate compartmentalization to achieve efficient and reversible immune inhibition, establishing LLPS as an intrinsic organizing principle of PD1 signaling.

## RESULTS

### PD1 microclusters are dynamic protein condensates

We first characterized the dynamics of PD1 microclusters in a cell–supported lipid bilayer (SLB) assay (*35*). We transduced PD1 and Shp2 double knockout (DKO) Jurkat cells with GFP-tagged human PD1 (*h*PD1) and mCherry-tagged human Shp2 (*h*Shp2), stimulated them with SLBs containing anti-*h*CD3ε antibody (α-*h*CD3ε), *h*ICAM-1, and *h*PDL1, and observed *h*PD1-GFP and mCherry-*h*Shp2 fluorescence using confocal fluorescence microscopy (**Fig. 1A**). As expected, *h*PD1 formed microclusters upon SLB contact and recruited *h*Shp2 to the plasma membrane (**Fig. 1B,C**). Time-lapse imaging revealed both fusion events of *h*PD1 microclusters, evidence of dynamic nature (**Fig. 1B**). Fluorescence recovery after photobleaching (FRAP) assay revealed a rapid photo-recovery of both *h*PD1 and *h*Shp2 within *h*PD1 microclusters (τ = 13.5 s for PD1, 8.3 s for Shp2, **Fig. 1D,E**), indicating dynamic exchange with the bulk phase. The slightly faster photo-recovery of *h*Shp2 compared to that of *h*PD1 might be a consequence of their distinct diffusion modes: on-membrane (2D) diffusion for *h*PD1 and in-solution (3D) diffusion for *h*Shp2. These data suggest that *h*PD1 microclusters have liquid-like properties.

**Fig. 1.**
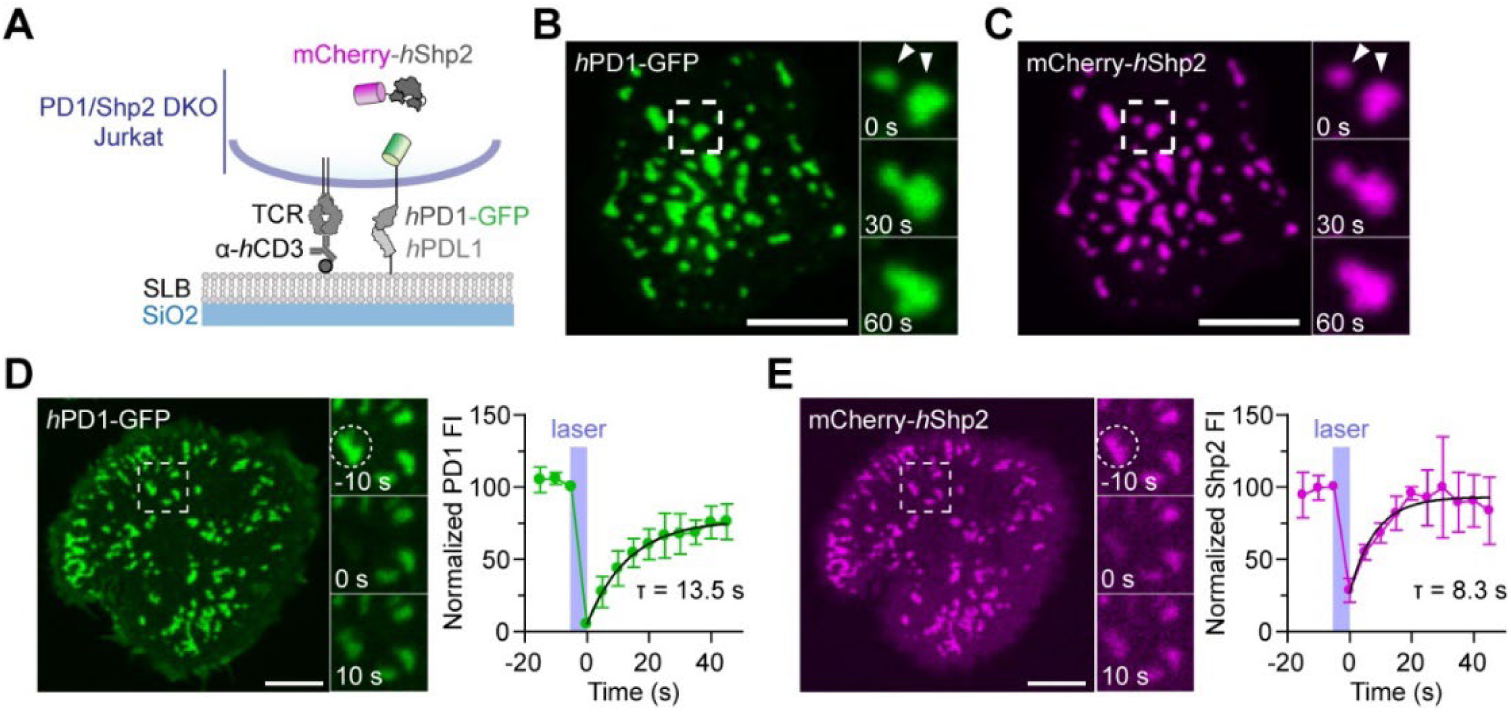
Dynamic nature of *h*PD1 microclusters in cells. **(A)** Schematic depicting the experimental set up for the cell–SLB assay, where a PD1/Shp2 DKO Jurkat cell expressing *h*PD1-GFP along with mCherry-*h*Shp2 contacts an SLB presenting α-*h*CD3ε (Okt3) and *h*PDL1. **(B)** Fusion events of *h*PD1 microclusters. Left, a representative confocal image of *h*PD1 microclusters in an SLB-bound Jurkat (*h*PD1-GFP) cell. Right, zoomed in time-lapse images of the boxed area shown on the left. Time zero indicates the beginning of image acquisition. **(C)** Left, a confocal image of mCherry-*h*Shp2 in the same cell as in B. Right, time-lapse images of the boxed area shown on the left. **(D)** FRAP of *h*PD1 microclusters. Left, a confocal image of *h*PD1 microclusters in an SLB-bound Jurkat (*h*PD1-GFP) cell. Middle, zoomed in time-lapse images of the boxed area of the left image, with the circled cluster photobleached at time zero. Right, time course of the fluorescent intensity (FI) of the circled cluster normalized to the FI of the adjacent unbleached cluster. n = 3 cells. **(E)** FRAP of mCherry-*h*Shp2 in the same cell as in D. n = 3 cells. Scale bars: 5 µm. Data are mean ± SD.

### ICD and pY motifs of hPD1 promote its clustering in stimulated T cells

We then examined the biochemical requirement for *h*PD1 clustering in a cell–SLB assay (**Fig. 2A**). We established three separate Jurkat T cell lines, each expressing similar levels of GFP-tagged *h*PD1 variants: WT (*h*PD1^WT^), a double tyrosine mutant (*h*PD1^FF^) in which tyrosine residues in ITIM and ITSM were both mutated to phenylalanine, or a truncation mutant lacking the ICD (*h*PD1^ΔICD^) (**Fig. 2B**). Confocal imaging showed that SLB-attached *h*PDL1 induced *h*PD1^WT^ clustering in Jurkat cells, consistent with previous reports (*10, 14*), but clustering decreased significantly for *h*PD1^FF^ and *h*PD1^ΔICD^ (**Fig. 2C**). Given that PD1^ICD^ associates with Shp2 through pY motifs, these data indicate that Shp2 binding contributes to *h*PD1 clustering.

**Fig. 2.**
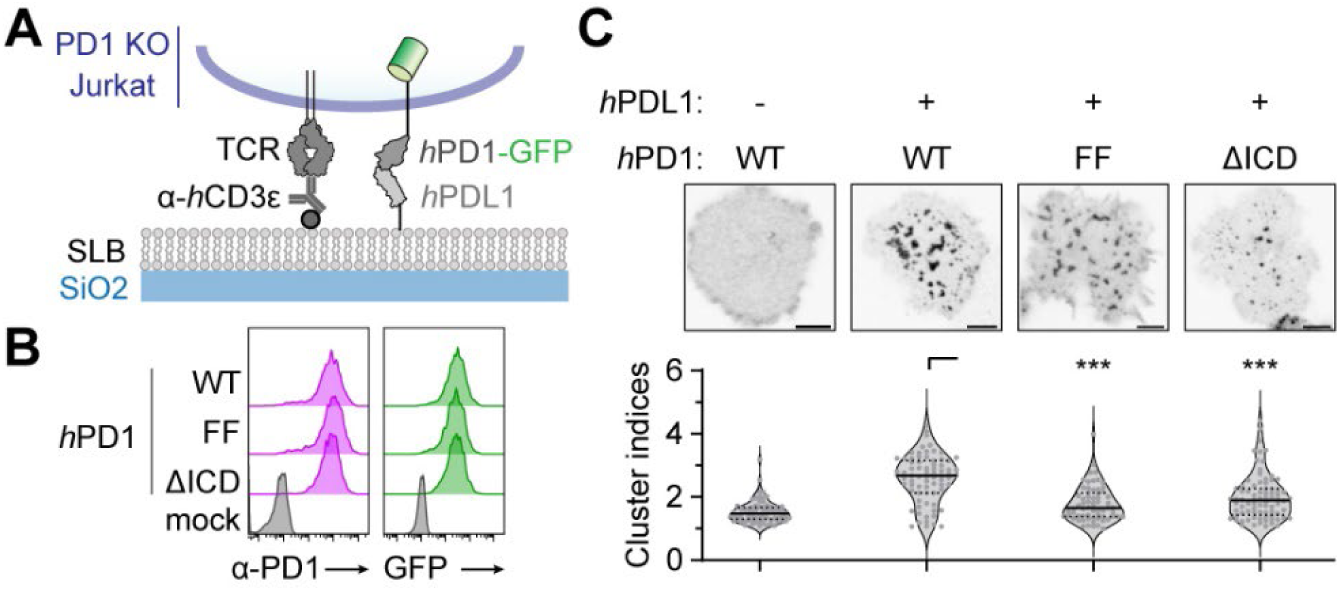
PD1 ICD and its pY motifs promote *h*PD1 microcluster formation. **(A)** Schematic depicting a PD1 KO Jurkat cell expressing GFP-tagged *h*PD1 stimulated on an SLB containing α-*h*CD3ε (Okt3) and *h*PDL1. **(B)** Flow cytometry showing the α-PD1 or GFP fluorescence of Jurkat cells expressing the indicated *h*PD1 variants. **(C)** Left, representative confocal images of GFP-tagged *h*PD1 variants in SLB-bound Jurkat cells as depicted in A. Right, *h*PD1 cluster indices under the indicated conditions. n = 58-79 cells. Scale bars: 5 µm. Data are mean ± SD. ***P < 0.001; student’s t-test.

### Purified Shp2 and phosphorylated PD1 co-formed liquid like condensates

We next investigated the mechanism by which *h*Shp2 binding promotes *h*PD1 microcluster formation using recombinant human proteins. We purified recombinant *h*Shp2^WT^, *h*PD1^ICD^, and phospho-*h*PD1^ICD^ (p-*h*PD1^ICD^) (**Fig. S1**) and mixed them in the presence of a crowding agent polyethylene glycol (PEG). Confocal imaging showed that a mixture of recombinant *h*Shp2^WT^ and phospho-*h*PD1^ICD^ (p-*h*PD1^ICD^) formed droplets that strongly enriched both proteins, accompanied by an increase in the turbidity of the solution, measured as optical density at 600 nm (OD_600_) (**Fig. 3A**, row 1). Neither protein alone formed droplets (**Fig. 3A**, rows 2 and 3), indicating that droplet formation depends on PD1:Shp2 interactions. Supportive of this notion, neither droplets nor turbidity increase was detected upon mixing *h*Shp2^WT^ with unphosphorylated *h*PD1^ICD^ (**Fig. 3A**, row 4). *h*Shp2^WT^:p-*h*PD1^ICD^ droplets underwent fusion events and recovered their fluorescence after photobleaching, indicating their liquid nature (**Fig. 3B,C**). These results demonstrate that PD1:Shp2 interactions can drive LLPS, potentially contribute to the observed PD1 clustering in cells.

**Fig. 3.**
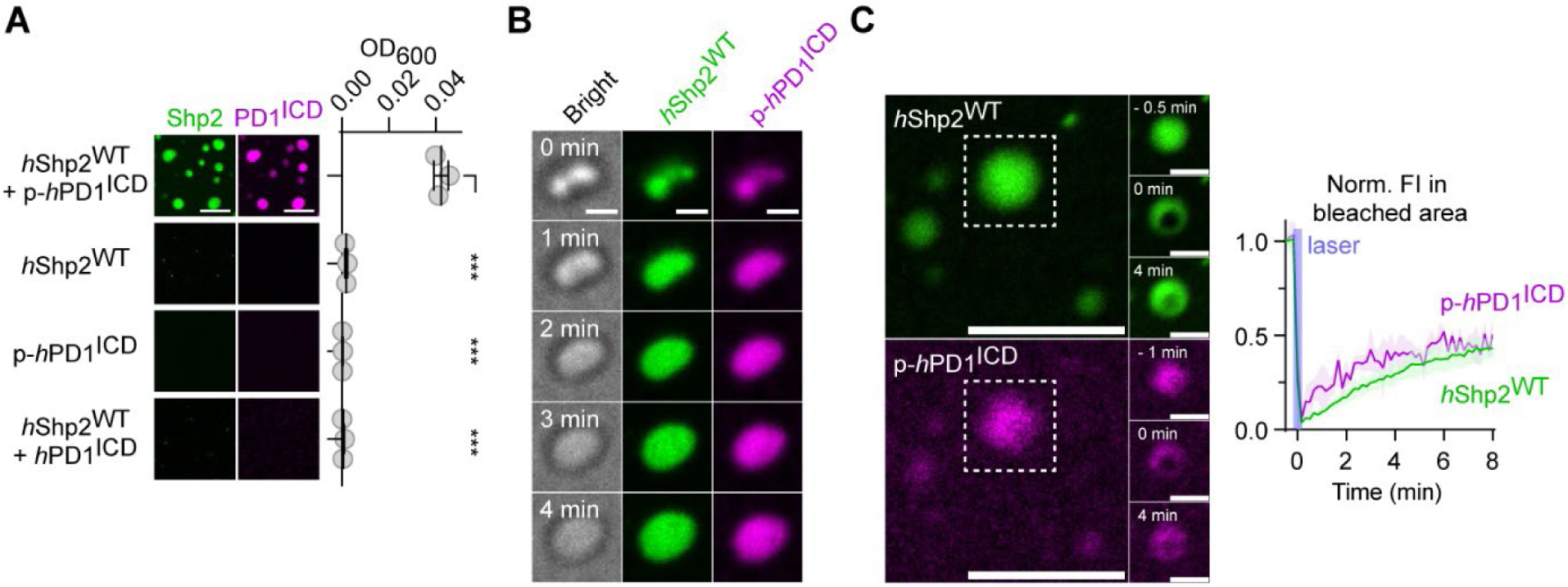
*h*PD1:*h*Shp2 interactions produce liquid-like condensates *in vitro*. **(A)** Left, confocal images of *h*Shp2^WT^ (green) mixed with the indicated *h*PD1^ICD^ (magenta) in LLPS buffer. Right, the OD_600_ values of the indicated *h*Shp2: *h*PD1^ICD^ mixture. [Shp2] = [PD1^ICD^] = 8 μM. n = 3 independent experiments. **(B)** Time-lapse bright field and confocal images showing a condensate fusion event in a mixture of 8 μM *h*Shp2^WT^ (green) and 8 μM p-*h*PD1^ICD^ (magenta). Time zero marks the start of image acquisition. **(C)** Left, time-lapse confocal images showing FRAP of *h*Shp2^WT^ (green) and *h*PD1^ICD^ (magenta) within the indicated condensate, which was photo-bleached at time zero. Right, FI time courses in the bleached area. FI values were normalized to those of a non-bleached *h*Shp2:p-*h*PD1^ICD^ condensate. [Shp2] = [pPD1^ICD^] = 8 μM. n = 3 condensates. Scale bars: 5 μm (A, C) or 1 μm (B, C inset). Data are mean ± SD. ***P < 0.001; student’s t-test.

### The catalytic activity of Shp2 confers the liquid property of PD1:Shp2 condensates

Shp2 may dephosphorylate PD1 to destabilize the PD1:Shp2 complex (*10, 11*). To examine how the catalytic activity of the PTPase domain affects PD1:Shp2 LLPS behavior, we compared full-length Shp2^WT^ with the catalytically dead Shp2 mutant: Shp2^C459E^, which naturally adopts a closed conformation, akin to Shp2^WT^ (*^36^*). Like the WT protein, *h*Shp2^C459E^ did not form condensates with non-phosphorylated *h*PD1^ICD^ but did so in the presence of p-*h*PD1^ICD^ (**Fig. 4A and Fig. S1**). However, FRAP analysis revealed that, unlike with the WT counterpart, *h*Shp2^C459E^:*h*PD1^ICD^ condensates failed to recover their fluorescence upon photobleaching (**Fig. 4B,C**), indicating their solid, gel-like nature. These results indicate that Shp2’s enzymatic activity confers the liquid nature of PD1:Shp2 condensates. By cleaving a subset of pY–SH2 interactions, Shp2 may destabilize the otherwise more static binding networks and allows molecules to rearrange and diffuse between the condensates and their surroundings.

**Fig. 4.**
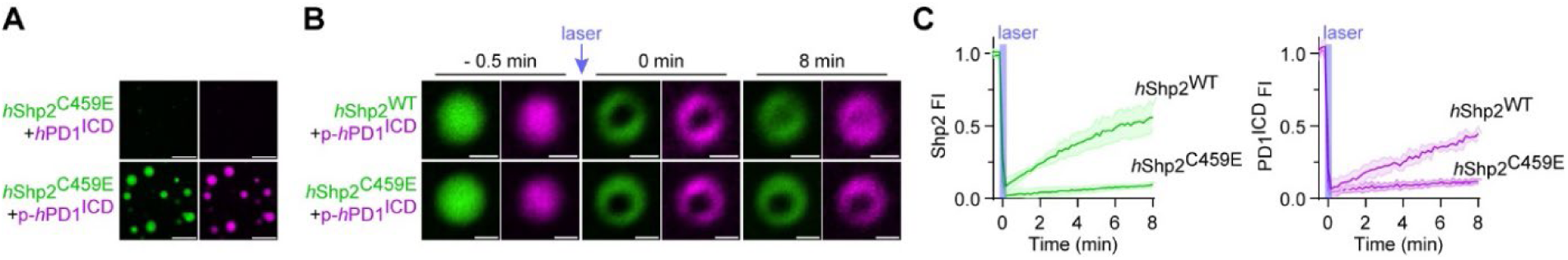
Shp2 catalytic activity promotes liquid-nature of PD1:Shp2 condensates. **(A)** Representative confocal images of *h*Shp2^C459E^ (green) mixed with either non-phosphorylated *h*PD1ICD or p-*h*PD1ICD (magenta). [Shp2] = [PD1^ICD^] = 8 μM. **(B)** Representative time-lapse confocal images of *h*Shp2:p-*h*PD1condensates containing the indicated Shp2 variant. Condensates were photo-bleached at time zero. [Shp2] = [PD1^ICD^] = 8 μM. **(C)** Time courses of Shp2 (left) or PD1^ICD^ (right) FI of the photo-bleached area in B. FIs were normalized to a non-bleached PD1:Shp2 condensates nearby. n = 3 condensates. Scale bars: 5 μm (A) or 1 μm (B). Data are mean ± SD.

### Electrostatic self-association of the Shp2 PTPase domain drives PD1:Shp2 LLPS

We next investigated the molecular basis of PD1:Shp2 LLPS. A previous study reported that the surface of Shp2’s PTPase domain contains acidic and basic surface patches that enable electrostatic self-association, driving LLPS of the isolated PTPase domain and of constitutively active Shp2 mutants that adopt an open conformation (*24*). In contrast, LLPS is restricted in the full-length form of Shp2, which normally exists in a closed conformation where the nSH2 covers the basic patch of the PTPase domain in *cis* (*24*). That study also showed that charge reversal mutations in the basic patch (R362E/K364E, abbreviated as REKE herein) prevent LLPS of Shp2 PTPase domain without affecting the intrinsic catalytic activity.

Because pY binding is known to convert Shp2 from a closed to an open conformation (*23, 25–28*), we hypothesized that PD1 binding exposes the basic patch on the Shp2 PTPase domain, thereby enabling Shp2 LLPS (**Fig. 5A**). Consistent with this notion, we found that increasing salt (NaCl) concentrations dose-dependently decreased p-*h*PD1:*h*Shp2 condensate formation (**Fig. 5B,C**) and dissolved the pre-formed condensates (**Fig. 5D**). Furthermore, both turbidity measurements and confocal imaging showed that the Shp2 REKE mutation markedly impaired the ability of full-length *h*Shp2 to form condensates with p-*h*PD1^ICD^ (**Fig. 5E and Fig. S1**). The same mutations similarly reduced the capacity of mouse Shp2 (*m*Shp2) to form condensates with p-*h*PD1^ICD^ (**Fig. 5F and Fig. S1**). These results indicate that pPD1:Shp2 LLPS is at least partly driven by electrostatic self-association of the Shp2 PTPase domain.

**Fig. 5.**
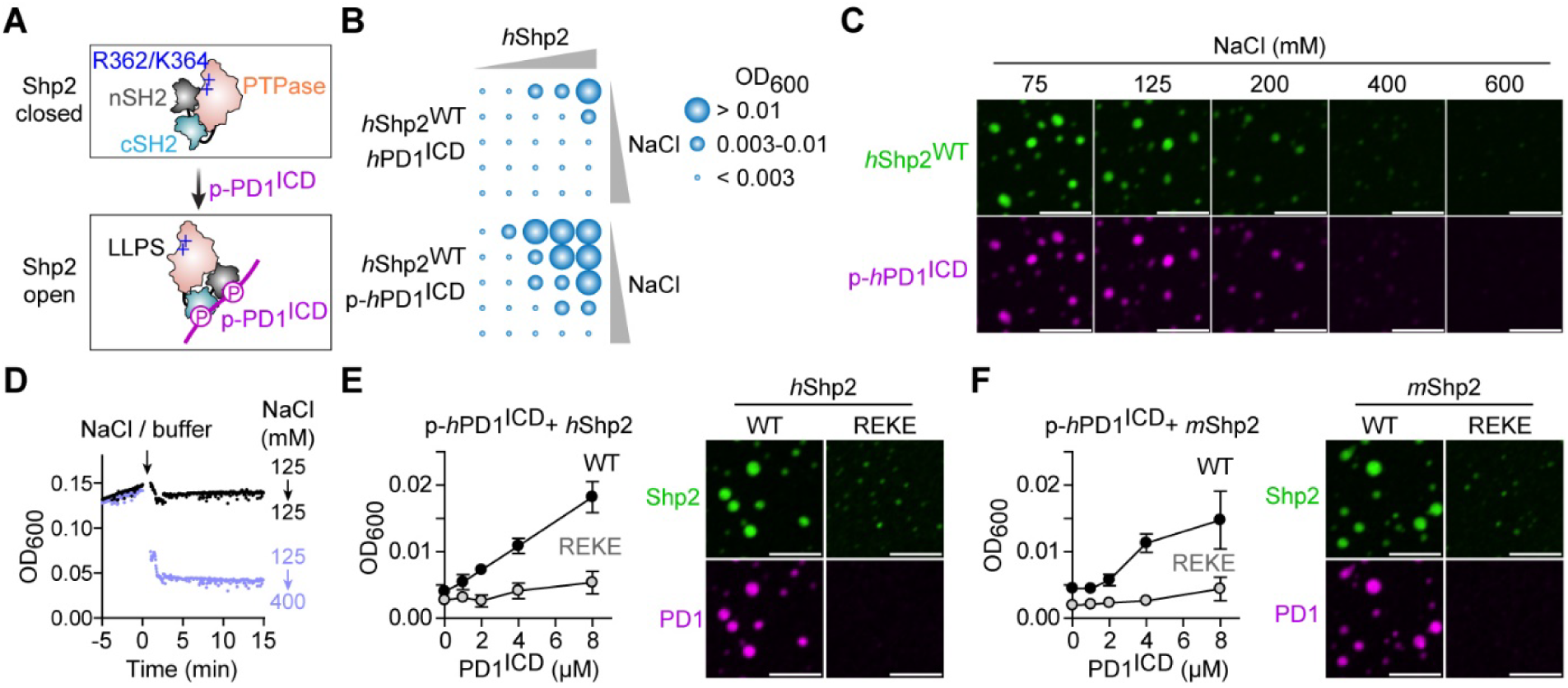
PD1:Shp2 LLPS is salt-sensitive and requires a basic patch in the Shp2 PTPase domain. **(A)** Cartoon depicting a model in which pPD1 induces a close-to-open conformational transition in Shp2. The basic amino acids that mediate Shp2 self-association (R362/K364) are highlighted as “+”. **(B)** Spot matrix plots showing the OD_600_ for *h*Shp2^WT^ mixed with either *h*PD1^ICD^ or p-*h*PD1^ICD^ across varying concentrations of Shp2 and NaCl. [PD1^ICD^] = 8 μM. From left to right, [Shp2] were 0, 2, 4, 8, 12 μM, from top to bottom [NaCl] were 75, 125, 200, 400, and 600 mM. **(C)** Confocal images of condensates formed by Shp2^WT^ and p-*h*PD1^ICD^ in LLPS buffer containing the indicated [NaCl]. **(D)** Time courses for the OD_600_ of Shp2^WT^:p-*h*PD1^ICD^ mixture before and after changes in [NaCl] upon injection of NaCl-containing LLPS buffer. [Shp2] = [PD1^ICD^] = 8 μM. **(E)** Effect of Shp2 REKE mutations on *h*PD1:*h*Shp2 LLPS. Left, OD_600_ of *h*Shp2^WT^ or *h*Shp2^REKE^ mixed with indicated [p-*h*PD1^ICD^]. Right, representative confocal images of the condensates in *h*PD1:*h*Shp2^WT^ mixture and *h*PD1:*h*Shp2^REKE^ mixture containing 8 μM of each. n = 3 independent experiments. **(F)** Same as E except that *m*Shp2 variants were used. n = 3 independent experiments. Scale bars: 5 μm. Data are mean ± SD.

### Ablation of PD1-induced Shp2 LLPS slowed CD3ζ dephosphorylation and reduced PD1–CD3ζ proximity

We then asked whether LLPS-disrupting mutations (REKE) alter Shp2 catalysis. Because Shp2 is autoinhibited, we compared *h*Shp2^WT^ and *h*Shp2^REKE^ in the presence of p-*h*PD1^ICD^ to bias *h*Shp2 toward an open, active conformation (*37*) (**Fig. 6A**). As expected, p-*h*PD1^ICD^ dose-dependently enhanced *h*Shp2^WT^-catalyzed turnover of the generic phosphatase substrate DiFMUP (**Fig. S2**). *h*Shp2^REKE^ dephosphorylated DiFMUP at rates similar to *h*Shp2^WT^, exhibiting a comparable *K*_M_ and a modestly higher *k*_cat_ (**Fig. 6B–D**). These assays were conducted under non-LLPS conditions, where *h*Shp2 and p-*h*PD1^ICD^ were maintained at nanomolar concentrations, well below the micromolar critical threshold. Thus, the REKE mutations do not impair the intrinsic catalytic activity of Shp2, agreeing with prior measurements using the isolated PTPase domain and an open-biased full-length Shp2 variant (*24*).

**Fig. 6.**
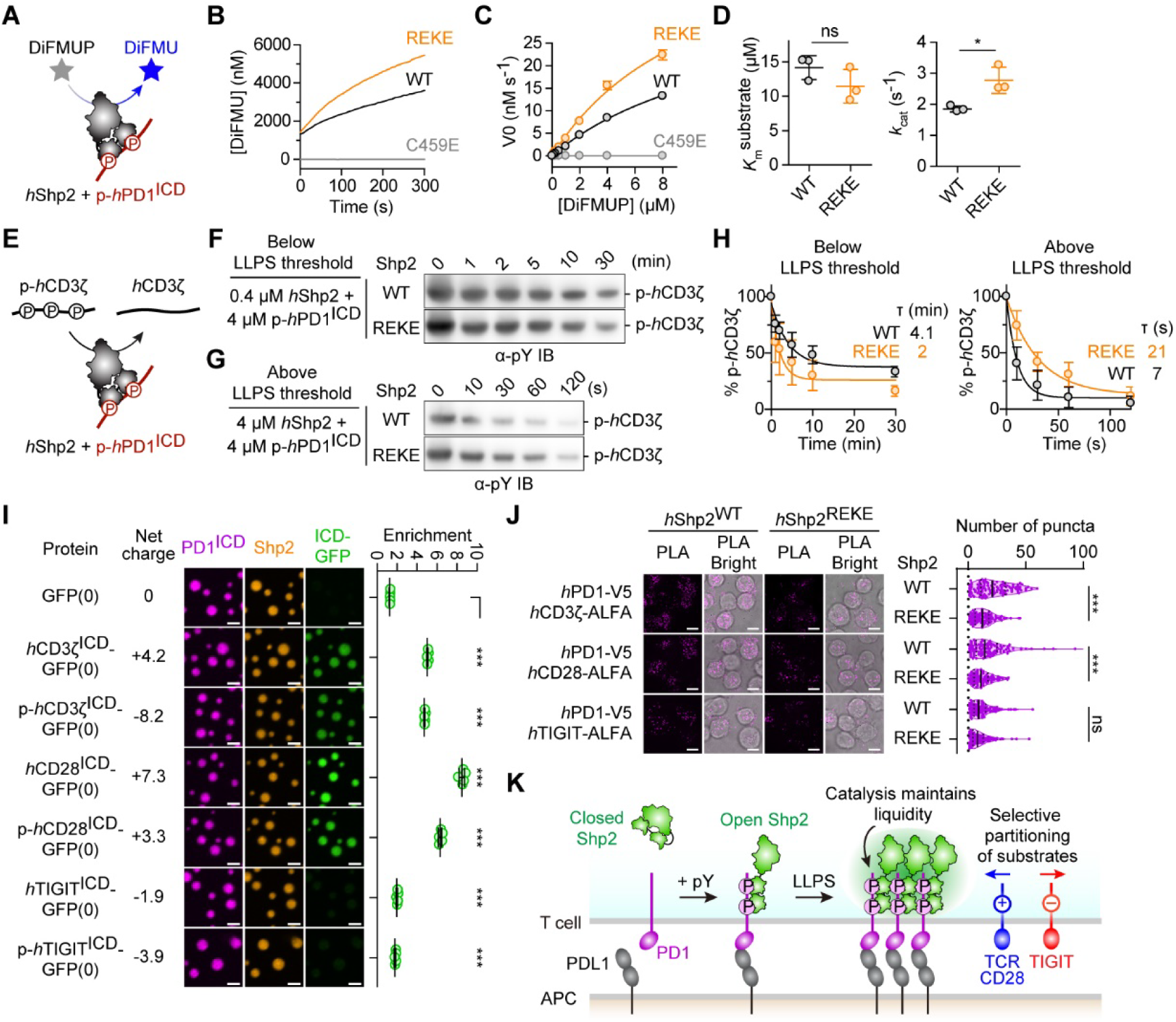
PD1:Shp2 LLPS accelerates substrate dephosphorylation by enriching targets within condensates. **(A)** Schematic depicting a Shp2 catalytic activity assay, where recombinant *h*Shp2 was incubated with p-*h*PD1^ICD^ to induce the open conformation. Shp2 mediated dephosphorylation of the substrate DiFMUP produces the fluorescent DiFMU. **(B)** Representative time course of [DiFMU] upon mixing with the indicated Shp2 variants. To prevent PD1:Shp2 LLPS, [*h*Shp2] and [p-*h*PD1^ICD^] were kept at 20 nM and 30 nM, respectively. **(C)** Initial velocity (V0) of [DiFMU] increase plotted against [DiFMUP] for the indicated Shp2 variants, fitted by the Michaelis-Menten model. n = 3 independent experiments. **(D)** Dot plots summarizing *K*_M_ and *k*_cat_ values calculated from C. n = 3 independent experiments. **(E)** Same as A, except that pre-phosphorylated *h*CD3ζ (p-*h*CD3ζ) was used as a substrate. **(F)** Representative IB showing the time-course of phosphorylation status of *h*CD3ζ upon mixing with 4 μM p- *h*PD1^ICD^ and 0.4 μM *h*Shp2^WT^ or *h*Shp2^REKE^. **(G)** Same as F, except that *h*CD3ζ was mixed with 4 μM p-*h*PD1^ICD^ and 4 μM *h*Shp2^WT^ or *h*Shp2^REKE^. **(H)** Time course of % p-*h*CD3ζ-GFP calculated from F (left) or G (right). n = 3 independent experiments. **(I)** Left, representative confocal images of *h*Shp2^WT^ (orange) and GFP(0)-tagged proteins (green) mixed with p-*h*PD1 (magenta) under the LLPS condition. Right, violin plots summarizing the GFP enrichment in the Shp2 condensates shown on the left. n = 5 independent experiments. **(J)** Left, representative confocal images of synaptic PLA signals (magenta) and bright field signals (grey) of Jurkat cells co-expressing V5-tagged *h*PD1 and the indicated ALFA-tagged receptors and *h*Shp2 variants. Right, violin plots summarizing the number of PLA puncta in individual cells shown on the left. n = 118-171 cells. **(K)** Model depicting how PD1:Shp2 LLPS organizes the inhibitory signalosome. PD1 phosphorylation promotes Shp2 condensation into dynamic LLPS compartments that enhance PD1 clustering, maintain liquidity through catalysis, and achieve selective substrate inclusion (CD3ζ/CD28) and exclusion (TIGIT). Scale bars: 2 μm (I) or 10 μm (J). Data are mean ± SD. *P < 0.05; ***P < 0.001; ns, not significant; student’s t-test.

We next examined a physiological substrate, CD3ζ, known as the principal signaling subunit of TCR (*38*) that recruit kinase Zap70 upon phosphorylation. Upon mixing p-*h*CD3ζ^ICD^, p-*h*PD1^ICD^, and *h*Shp2^WT^ or *h*Shp2^REKE^, under non-LLPS conditions, α-pY immunoblots (IBs) showed that p-*h*CD3ζ^ICD^ was dephosphorylated more rapidly by *h*Shp2^REKE^ (τ = 2 min) than by *h*Shp2^WT^ (τ = 4.1 min) (**Fig. 6E,F,H**), consistent with the notion that Shp2 REKE mutant preserves (or modestly enhances) intrinsic PTPase activity. By contrast, under LLPS conditions in which Shp2 concentration was increased to 4 μM, *h*Shp2^REKE^-mediated dephosphorylation was three-fold slower than that of *h*Shp2^WT^ (τ = 21 s for *h*Shp2^REKE^ versus τ = 7 s for *h*Shp2^WT^) (**Fig. 6G,H**). These data indicate that PD1:Shp2 LLPS accelerates CD3 dephosphorylation.

Both *h*PD1^ICD^ and *h*Shp2 carry net negative charges at pH 7.4 (*h*PD1^ICD^ = – 2.0; *h*Shp2 = – 3.2) based on their predicted isoelectric points, indicating that PD1:Shp2 LLPS are overall negatively charged. We therefore hypothesized that PD1-induced Shp2 LLPS promotes substrate co-compartmentalization through electrostatic interactions. To test this, we pre-formed *h*Shp2^WT^:p-*h*PD1^ICD^ condensates, and mixed them with either WT GFP (net -7 charge) or its charge-modified mutant GFP(0) (neutral), or GFP(+7) (net +7 charge). Confocal imaging showed that PD1:Shp2 condensates selectively incorporated GFP(+7), but not GFP WT or GFP (0), indicating a preference for positively charged molecules (**Fig. S3**). Next, we purified GFP(0)-tagged ICDs of CD3ζ, the costimulatory receptor CD28, the coinhibitory receptor TIGIT in both phosphorylated and unphosphorylated forms (see Methods), and mixed them with pre-formed *h*Shp2^WT^:p-*h*PD1^ICD^ condensates. We observed pronounced enrichment of CD3ζ^ICD^ and CD28^ICD^ within PD1:Shp2 condensates compared to TIGIT^ICD^ (**Fig. 6I**). These experiments revealed that PD1:Shp2 condensates enrich positively charged protein clients, with the exception of p-CD3ζ^ICD^, which was strongly enriched despite carrying a net charge of – 8.2, indicating that factors other than net charge also contribute to the molecular partition. These results suggest that Shp2 LLPS assembles a PD1 signalosome that selectively concentrates positively charged substrates/co-receptors, akin to charge-governed partitioning in the LAT signalosome (*17*).

To determine whether the molecular enrichment of PD1:Shp2 condensates translates to molecular proximity in cells, we performed proximity ligation assays (PLA) (*39, 40*) in Jurkat cells expressing V5-tagged *h*PD1 together with ALFA-tagged human CD3ζ, CD28, or TIGIT, in the presence of either *h*Shp2^WT^ or *h*Shp2^REKE^ (**Fig. S4**). Cells were stimulated on SLBs bearing *h*PD-L1 and α-*h*CD3ε alone or along with human CD86 (ligand for CD28) or PVR (ligand for TIGIT). Synaptic PLA signal visualized by confocal microscopy revealed greater PD1 proximity to CD3ζ and CD28 in Shp2^WT^-expressing cells compared with Shp2^REKE^-expressing cells, whereas PD1:TIGIT proximity was considerably weaker and unaffected by REKE mutations (**Fig. 6J**).

Collectively, these data support the notion that PD1-induced Shp2 LLPS compartmentalizes PD1 with its substrates, based on electric charges and additional mechanisms, enabling efficient inhibitory signaling (**Fig. 6K**).

### Disruption of PD1-induced Shp2 LLPS impairs PD1 clustering and inhibitory function

Having established that PD1-induced Shp2 LLPS promotes the colocalization of PD1 with its signaling substrates and accelerates CD3ζ dephosphorylation (**Fig. 6**), we next asked whether this LLPS process operates in cells to promote PD1 microcluster formation and inhibitory signaling. To address this, we leveraged the LLPS-disrupting Shp2 REKE mutant in cell-based imaging and functional assays.

We examined whether disrupting Shp2 LLPS alters PD1 microcluster formation in cells in the established cell–SLB assay with TIRF imaging. In *h*PD1-GFP transduced, *h*Shp2 KO Jurkat cells, we lentivirally expressed either mCherry-*h*Shp2^WT^ or mCherry-*h*Shp2^REKE^ at comparable levels (**Fig. 7A**). Upon contacting PDL1/αCD3ε SLBs, cells expressing *h*Shp2^WT^ formed more intense PD1 microclusters that recruited more Shp2 than those expressing *h*Shp2^REKE^ (**Fig. 7B**). These results indicate that PD1:Shp2 LLPS facilitates the co-enrichment of PD1 and Shp2 within signaling clusters.

**Fig. 7.**
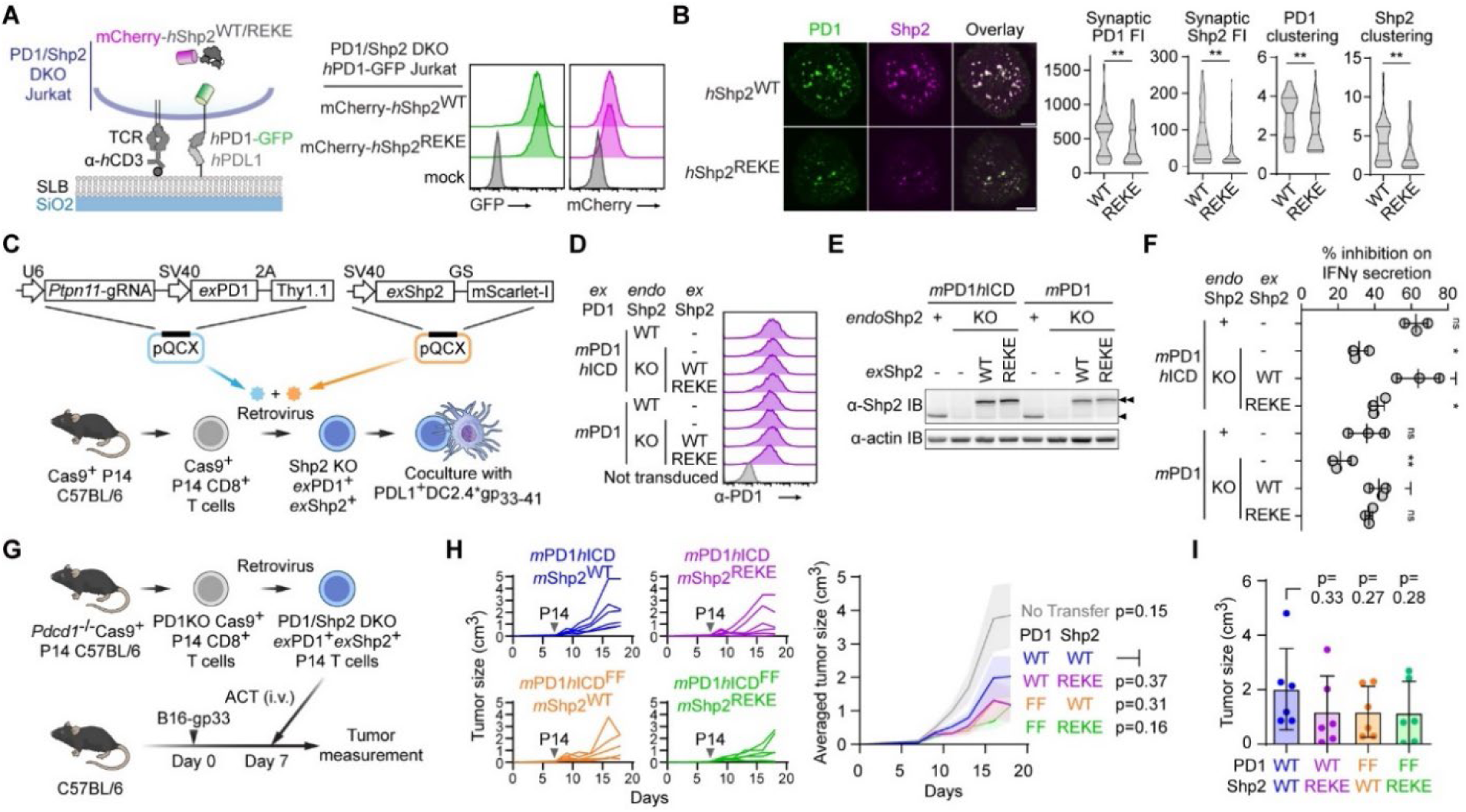
Shp2 LLPS promotes the microcluster formation and inhibitory function of PD1. **(A)** Left, schematic depicting a Jurkat cell expressing *h*PD1-GFP along with either mCherry-tagged *h*Shp2^WT^ or *h*Shp2^REKE^, contacting an SLB presenting Okt3 and *h*PDL1. Right, flow cytometry showing GFP and mCherry fluorescent intensities of the indicated Jurkat cells. **(B)** Left, representative confocal images showing PD1 (green) and Shp2 (magenta) at the interface of Jurkat cells and SLBs shown in A. Right, violin plots showing the synaptic fluorescent intensities or clustering indices of PD1 or Shp2. **(C)** Schematic depicting a P14:DC2.4 coculture assay. Cas9^+^ P14 cells were retrovirally transduced with Shp2-targeting gRNA, along with exogenous Shp2 (*ex*Shp2: mScarlet-*m*Shp2^WT^ or *m*Shp2^REKE^) and exogenous PD1 (*ex*PD1: *m*PD1^WT^ or *m*PD1*h*ICD), and cocultured with gp_33-41_ peptide-pulsed PDL1^+^ DC2.4 cells. **(D)** Flow cytometry showing the PD1 expression on the indicated P14 cells prepared in C. **(E)** IBs showing the *endo*Shp2 or *ex*Shp2 expression in the indicated P14 cells prepared in C. **(F)** % inhibition on IFNγ secretion under the indicated conditions calculated from Fig. S6. **(G)** Schematic depicting an ACT murine melanoma model. PD1 KO Cas9^+^ P14 cells were retrovirally transduced with *Ptpn11*-targeting gRNA, alongside *ex*PD1 (*m*PD1*h*ICD or *m*PD1*h*ICD^FF^) and *ex*Shp2 (mScarlet-*m*Shp2^WT^ or *m*Shp2^REKE^), and intravenously injected (i.v.) into host mice bearing B16-gp33 melanoma. n = 6–8 mice. **(H)** Tumor growth curves in mice that received P14 cells expressing the indicated *ex*PD1 and *ex*Shp2. **(I)** Endpoint tumor sizes in H. Scale bars: 5 μm. Data are mean ± SD. *P < 0.05; **P < 0.01; ns, not significant; student’s t-test (B, F, and I) or two-way ANOVA (H).

We next asked whether the impaired clustering caused by the Shp2 REKE mutation translates into reduced PD1 inhibitory signaling. We measured secreted IL-2 from the aforementioned Jurkat cells following stimulation with *h*PDL1-expressing Raji cells pulsed with superantigen staphylococcal enterotoxin E (SEE), in the presence or absence of α-PDL1-blockade antibody atezolizumab (**Fig. S5A**). In *h*Shp2^WT^-expressing cells, α-PDL1 approximately doubled IL-2 production, indicating ∼50% inhibition of T cell activity by PD1 (**Fig. S5B**). By contrast, the effect of α-PDL1 was weaker in *h*Shp2^REKE^-expressing cells, corresponding to ∼26% inhibition (**Fig. S5B**), consistent with reduced PD1 clustering and Shp2 recruitment. Interestingly, IL-2 secretion was higher in *h*Shp2^REKE^-expressing cells even under PD1-blocking conditions, suggesting that Shp2 LLPS may also regulate ligand-independent tonic PD1 signaling (*41*) or TCR signaling.

To further test this mechanism in primary T cells, we isolated Cas9^+^ P14 CD8^+^ mouse T cells, which recognize the H-2D^b^ (MHC-I)-restricted lymphocytic choriomeningitis virus (LCMV) peptide antigen gp_33-41_. We generated four groups of P14 cells by co-transducing with two retroviruses. The first encoded a gRNA targeting *Ptpn11* (*m*Shp2) together with either *m*PD1 WT or *m*PD1 with its ICD humanized (*m*PD1*h*ICD). The second encoded an mScarlet-tagged exogenous Shp2 variant (*ex*Shp2^WT^ or *ex*Shp2^REKE^). This co-transduction strategy enabled replacement of *endo*Shp2 with defined *ex*Shp2 variants, while simultaneously reconstituting specific PD1 variants (**Fig. 7C**). Control P14 groups expressed *m*PD1 variants along with either a Shp2-targeting gRNA (Shp2KO) or a non-targeting gRNA (*endo*Shp2). Flow cytometry confirmed comparable PD1 expression across all conditions (**Fig. 7D**), and immunoblotting verified efficient deletion of endogenous *m*Shp2 and similar expression of mScarlet-tagged *ex*Shp2 variants (**Fig. 7E**).

We then quantified secreted IFNγ after co-culture of P14 cells with PDL1⁺ DC2.4 cells pulsed with gp_33-41_ peptide, in the presence or absence of α-PD1 blockade antibody. PD1 blockade increased IFNγ secretion for all P14 cells, confirming the PD1-mediated inhibitory function (**Fig. S6**). Shp2 KO reduced the inhibitory functions of *m*PD1*h*ICD and *m*PD1 by 50% and 40%, respectively (**Fig. 7F**). Reconstitution with *ex*Shp2^WT^ restored the inhibitory activity of both PD1 variants, whereas *ex*Shp2^REKE^ only partially rescued the function of *m*PD1*h*ICD. Based on these comparisons, Shp2 REKE mutation attenuated *m*PD1*h*ICD function by ∼69% and *m*PD1 function by ∼26% (**Fig. 7F**), suggesting a greater contribution of Shp2 LLPS to human PD1 signaling that is consistent with the weaker ability of *m*PD1 to bind Shp2 (*42*) and to induce Shp2 LLPS (**Fig. S1 and S7**). Flow-cytometric analysis of intracellular cytokine staining yielded similar trends: Shp2 REKE mutation attenuated *m*PD1*h*ICD function by ∼73% and *m*PD1 function by ∼40% (**Fig. S8**).

Finally, we examined whether Shp2 LLPS contributes to PD1 inhibitory function *in vivo* using an adoptive T cell transfer (ACT) murine melanoma model (**Fig. 7G**). PD1–deficient P14 CD8^+^ T cells were reconstituted with either WT PD1 ICD (*m*PD1*h*ICD) or a signaling-defective tyrosine-motif mutant (*m*PD1*h*ICD^FF^), together with either WT Shp2 or an LLPS-deficient Shp2 mutant (REKE), and transferred into C57BL/6 mice bearing gp_33-41_-expressing B16 melanoma (B16-gp33). Across groups, *ex*PD1 and *ex*Shp2 were expressed at comparable levels (**Fig. S9**). As expected, P14 transfer slowed tumor growth relative to no transfer (**Fig. 7H**). Within the *m*PD1*h*ICD background, Shp2^WT^ cells trended toward weaker tumor control than Shp2^REKE^ cells (**Fig. 7H and I**). In contrast, under the *m*PD1*h*ICD^FF^ background, P14 cells expressing *ex*Shp2^WT^ or *ex*Shp2^REKE^ exhibited comparable anti-tumor ability (**Fig. 7H and I**). These data are consistent with a model in which Shp2 LLPS supports PD1–dependent inhibition in vivo, and suggest that disrupting Shp2 LLPS as a strategy to weaken PD1 function while boosting T cell anti-tumor immunity.

## DISCUSSION

This study identifies LLPS of Shp2 as a central mechanism that drives PD1 clustering, substrate organization, and inhibitory signaling. PD1 engagement with its ligand PDL1 licenses Shp2 self-association, producing dynamic PD1:Shp2 condensates whose liquid properties depend on Shp2 catalytic activity. Disruption of this self-association weakens PD1 microcluster formation, substrate colocalization, and inhibitory function, establishing LLPS as an intrinsic organizing principle of the PD1 pathway.

PD1 has long been known to form microclusters upon ligand engagement (*10–12*), yet the origin and purpose of these structures have remained unclear. Our data show that these clusters behave as dynamic condensates whose formation arises from multivalent pY:SH2 interactions and electrostatic self-association of Shp2 via its PTPase domain. Shp2 LLPS might synergize with PD1 transmembrane-domain dimerization, as reported recently (*43*), to promote PD1 clustering. LLPS thereby links receptor phosphorylation, Shp2 activation, and molecular compartmentalization, converting discrete molecular interactions into a reversible inhibitory hub that enriches phosphatase components while maintaining rapid cytoplasmic exchange.

We provided evidence that PD1:Shp2 LLPS can serve to organize signaling molecules on the basis of electric charges, with positively charged CD3ζ and CD28 ICDs selectively enriched over negatively charged TIGIT ICD. Indeed, positively charged patches appear to be a common property of the ICDs of stimulatory receptors (*44–46*). This electrostatic sorting accelerates dephosphorylation of key substrates such as CD3ζ, thereby enhancing signal attenuation. Conversely, reduced partitioning of TIGIT ICD suggests that PD1 condensates may help maintain pathway specificity by preferentially recruiting activating substrates while limiting access of certain inhibitory receptors. LLPS thus creates an electrostatically tuned microenvironment that promotes pathway-specific inhibition, reminiscent of charge-based partitioning in the LAT signalosome (*17*). This does not exclude other mechanisms, indeed, the phosphorylated form of CD3ζ^ICD^, which carries a net negative charge, still partitioned into Shp2 condensates, indicating that other biochemical properties, including the patterning of charges (*47*), also contribute to the molecular partitioning.

The dependence of condensate liquidity on Shp2 enzymatic turnover suggests a self-regulating feedback mechanism, where dephosphorylation of pY:SH2 bonds remodels the condensate network and sustains fluidity, analogous to ATP-driven remodeling in protein:RNA condensates (*48–50*). This coupling between catalysis and material state exemplifies how energy-dependent enzymatic cycles maintain dynamic signaling assemblies.

The electrostatic interface within the Shp2 PTPase domain is pivotal for LLPS. Charge-reversal mutations (REKE) abolished condensation, reduced substrate co-partitioning, and attenuated PD1 function, indicating that reversible self-association is integral to both spatial organization and signaling output. This same surface drives constitutive LLPS in oncogenic Shp2 mutants (*24*), implying that normal and pathological condensates share a physical basis but differ in regulation. In WT Shp2, LLPS is phosphorylation-dependent; in pathological mutants, it becomes unrestrained.

We further found that human PD1 induced Shp2 condensation more efficiently than its murine counterpart, consistent with stronger PD1:Shp2 binding in humans (*42*). This species divergence highlights evolutionary tuning of PD1:Shp2 coupling and may contribute to differences in immune checkpoint sensitivity between humans and mice.

Functionally, PD1-induced Shp2 condensates act as liquid-like, transient inhibitory hubs that concentrate phosphatase activity near TCR and CD28 complexes, enabling potent yet reversible suppression of T cell activation. By assembling and dissolving in a signal-dependent manner, PD1–induced Shp2 condensates can deliver localized, tunable, and reversible inhibition.

In summary, PD1 engagement triggers catalytically regulated phase transition of Shp2 that couples enzymatic activation, substrate compartmentalization, and mesoscale assembly to enforce inhibitory signaling. These findings position PD1 within the emerging paradigm of phase-separated signalosomes and suggests new strategies to tune immune inhibition by targeting the material or electrostatic properties of checkpoint condensates.

## Acknowledgments

We thank E. Griffis and P. Guo at the Nikon Imaging Center of UCSD for technical assistance in confocal imaging and K. Khow and J. Zhang for critically reading the manuscript. T.M. was supported by a Human Frontiers Science Program Long-Term Fellowship (LT000909/2019-L) and JST, PRESTO (JPMJPR22EB). This work was supported by National Institute of Health grant R37 CA239072 (to E.H.), American Cancer Society grants AWD103481 and CAT-24-1374685-01-CAT (to E.H.); and a Department of Defense award HT9425-25-1-0929 (to E.H.).

## Author contributions

T.M., Conceptualization, Data curation, Formal analysis, Funding acquisition, Investigation, Methodology, Project administration, Resources, Software, Validation, Visualization, Writing – original draft, Writing – review & editing; G.A.W., Investigation, Methodology, Resources, Validation, Writing – review & editing; X.S., Resources, Writing – review & editing; K.G., Investigation, Resources; H.S., Resources, Writing – review & editing; C.W., Resources, Writing – review & editing; E.H., Conceptualization, Funding acquisition, Methodology, Project administration, Supervision, Visualization, Writing – original draft, Writing – review & editing.

## Data and materials availability

All data are available in the main manuscript or the supplementary materials. Requests for materials should be addressed to E.H.

---

**Fig. S1.**
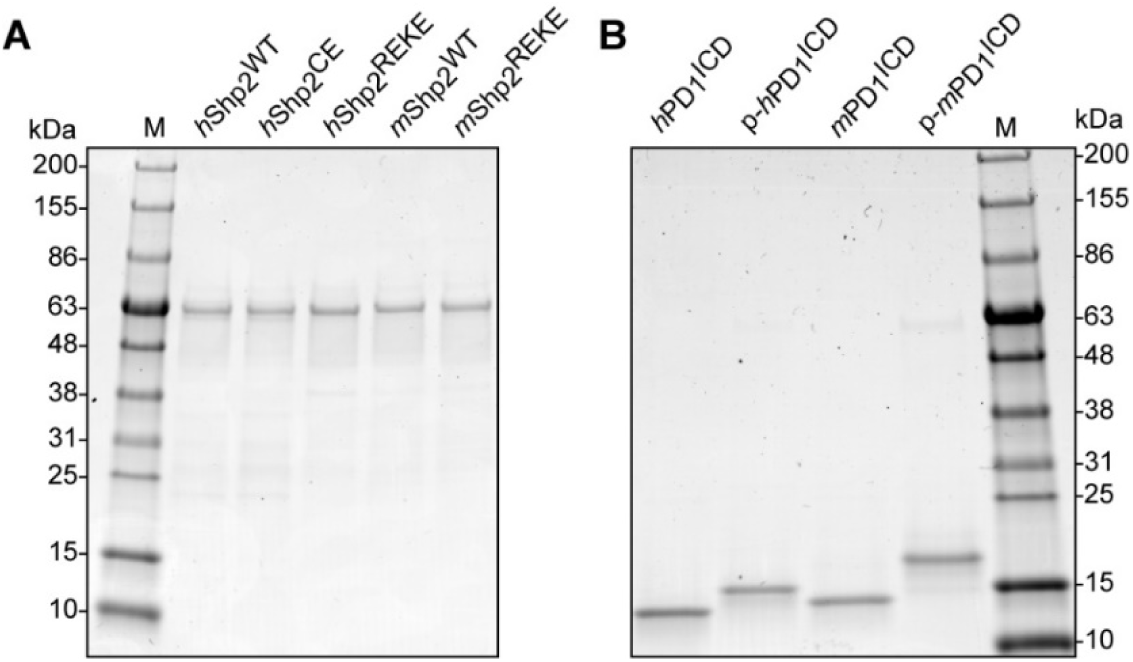
Coomassie-stained SDS-PAGE of purified proteins used in this study. **(A)** Coomassie-stained SDS-PAGE of 0.5 μg of purified Shp2 variants **(B)** Coomassie-stained SDS-PAGE of 0.5 μg of purified PD1 variants.

**Fig. S2.**
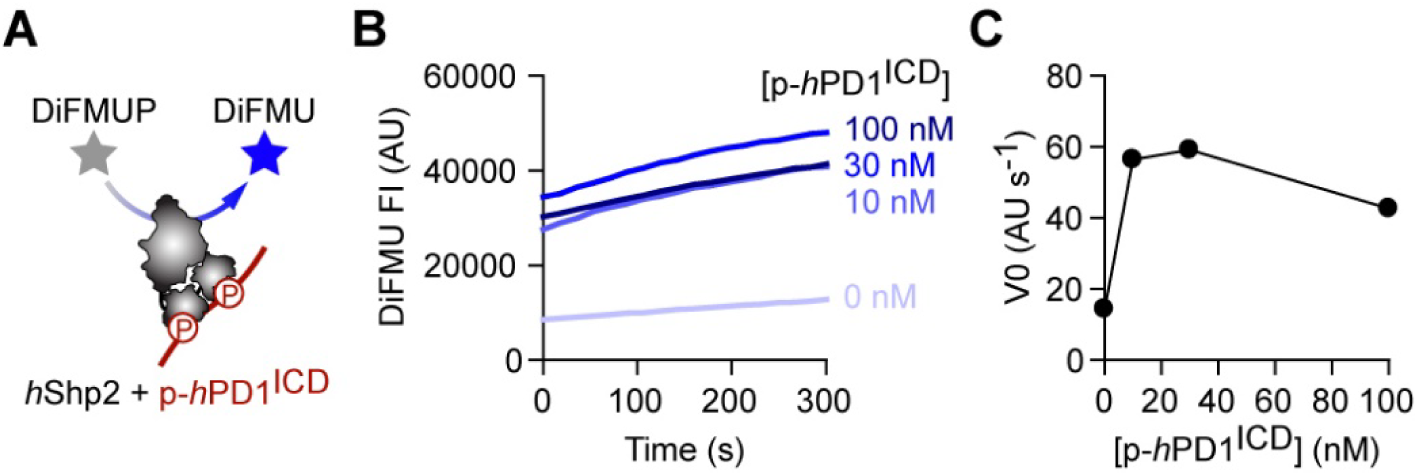
p-PD1^ICD^ potentiates Shp2 catalytic activity *in vitro*. **(A)** Schematic depicting a Shp2 catalytic activity assay, where recombinant *h*Shp2 was incubated with p-*h*PD1^ICD^ to induce the open conformation. Shp2 mediated dephosphorylation of the substrate DiFMUP produces the fluorescent DiFMU. **(B)** Representative time course of DiFMU FI upon mixing with 20 nM *h*Shp2^WT^ and the varying concentrations of [p-*h*PD1^ICD^]. **(C)** Initial velocity (V0) of DiFMU FI increase plotted against [p-*h*PD1^ICD^] calculated from B.

**Fig. S3.**
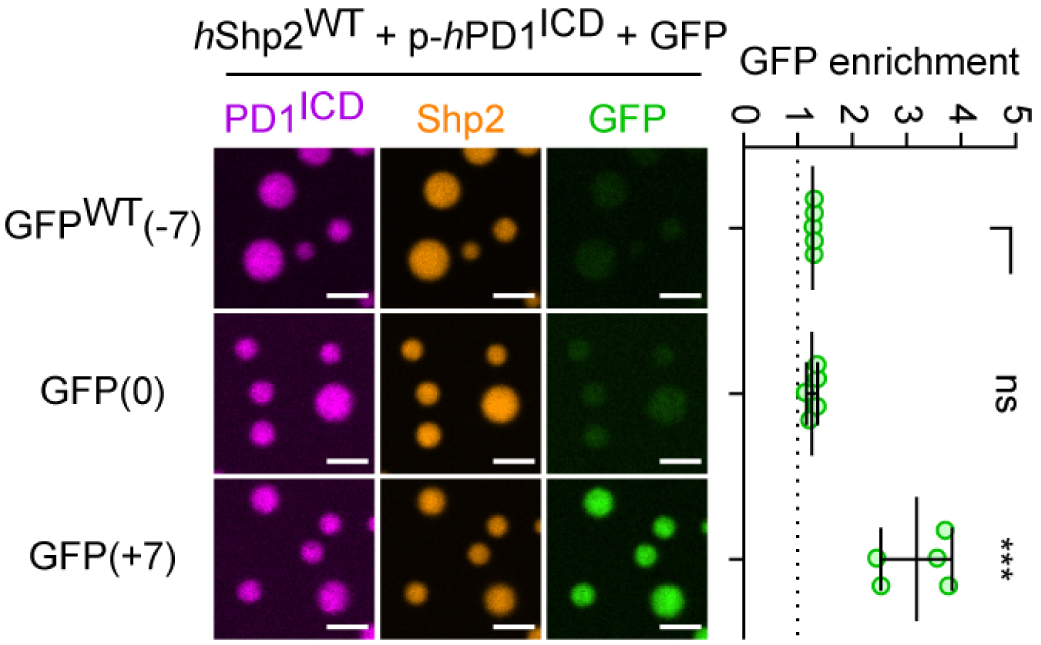
Charge-dependent guest protein partitioning into PD1:Shp2 LLPS. Left, representative confocal images of the mixture of p-*h*PD1 (magenta), *h*Shp2^WT^ (orange), and the indicated charge-modified GFP (green). Right, dot plots summarizing the GFP enrichment in the PD1:Shp2 condensates shown on the left. n = 5 independent experiments. Scale bars: 2 μm. Data are mean ± SD. ***P < 0.001; ns, not significant; student’s t-test.

**Fig. S4.**
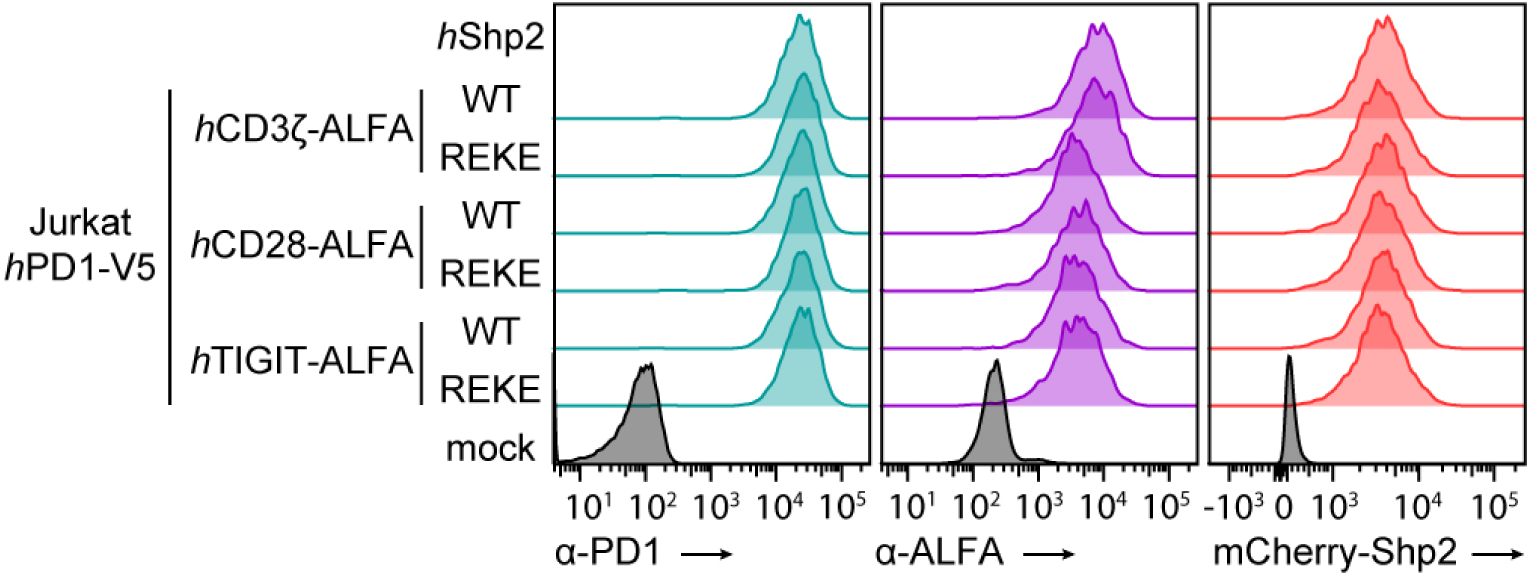
Relevant protein expressions in cell lines used for the PLA-SLB assay. Flow cytometry showing the expression of V5-tagged *h*PD1 (left), ALFA-tagged receptors (middle), and mCherry-tagged *h*Shp2 (right) in the indicated Jurkat cells.

**Fig. S5.**
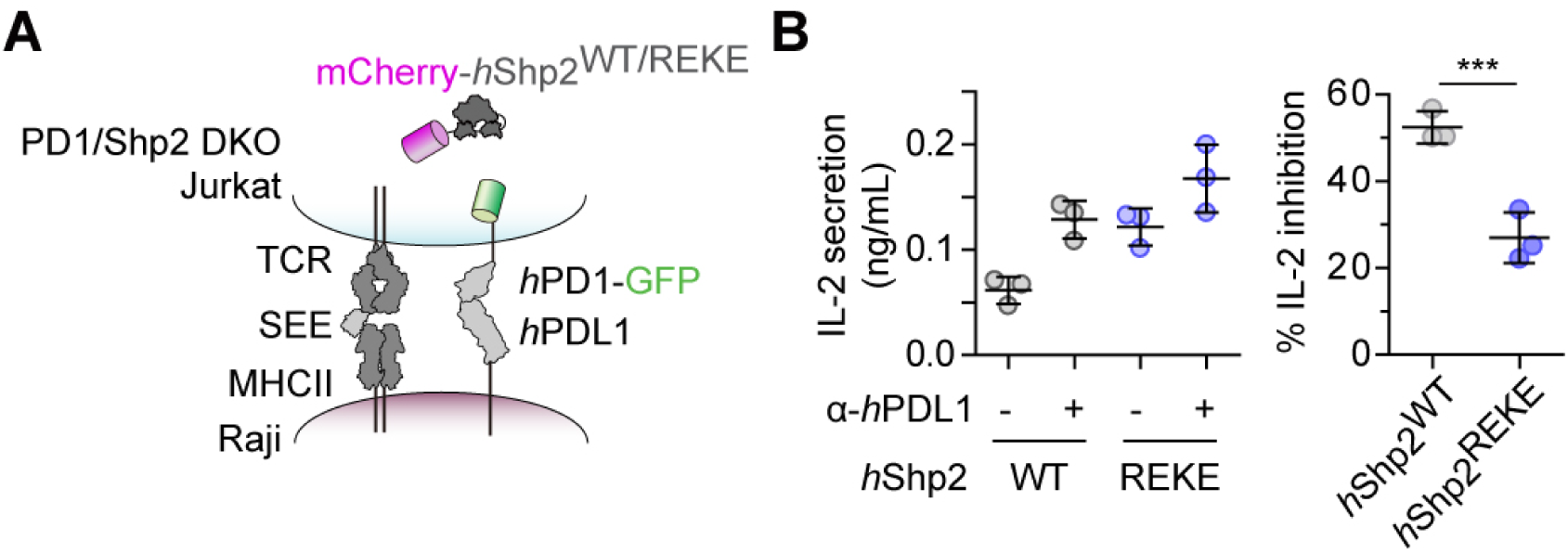
Effects of Shp2 REKE mutations on PD1 inhibitory function in Jurkat cells. **(A)** Schematic depicting a Jurkat cell expressing *h*PD1-GFP with either mCherry-*h*Shp2^WT^ or *h*Shp2^REKE^, cocultured with SEE-loaded, *h*PDL1-expressing, CD80 KO Raji cells. **(B)** Left, IL-2 secreted in the Jurkat:Raji coculture shown in a in the presence or absence of α-*h*PDL1. Right, % inhibition of IL-2 secretion mediated by the indicated *h*PD1 variant calculated from the left. [SEE] = 0.06 ng/mL. n = 3 independent experiments. Data are mean ± SD. ***P < 0.001; student’s t-test.

**Fig. S6.**
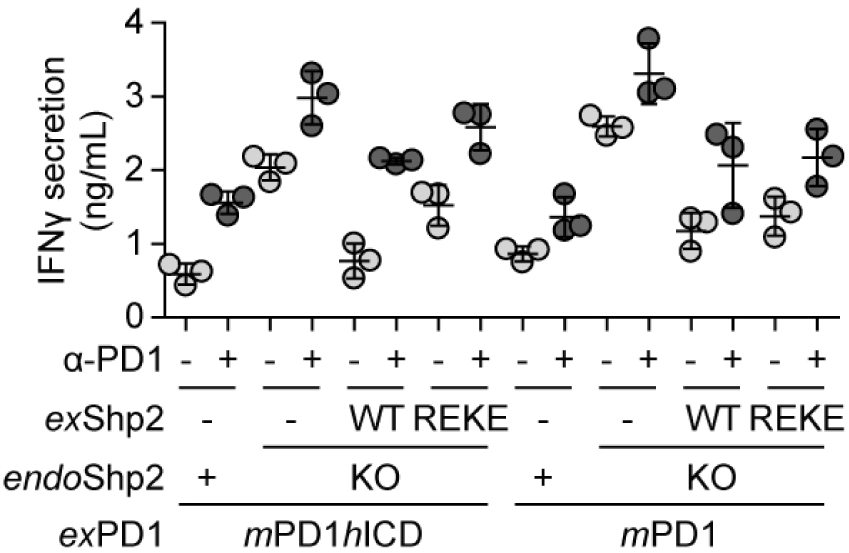
IFNγ secretion from P14 cells expressing the indicated genes in the presence or absence of α-PD1 blockade. Dot plots of secreted IFNγ in P14: DC2.4 cocultures shown in Fig. 7C, measured by ELISA.

**Fig. S7:**
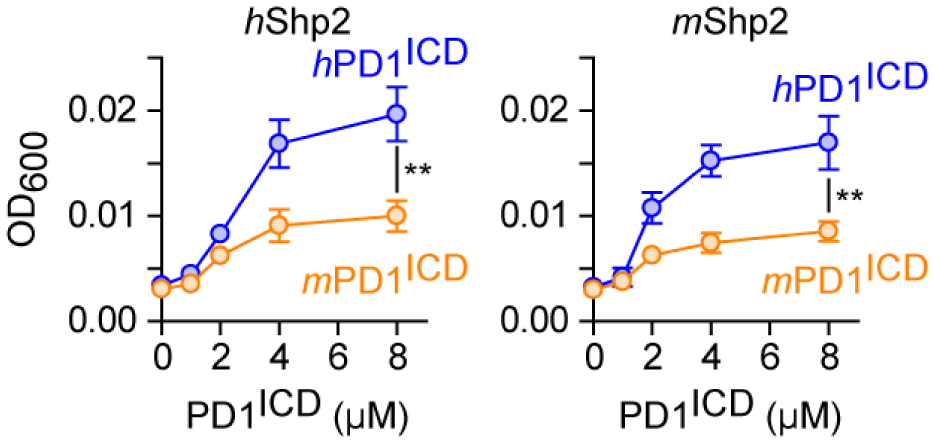
hPD1 more strongly induces Shp2 LLPS than does *m*PD1. OD_600_ of 8 μM *h*Shp2^WT^ or *m*Shp2^WT^ mixed with either pre-phosphorylated *h*PD1^ICD^ or *m*PD1^ICD^ at the indicated concentration. n = 3 independent experiments. Data are mean ± SD. **P < 0.01; student’s t-test.

**Fig. S8.**
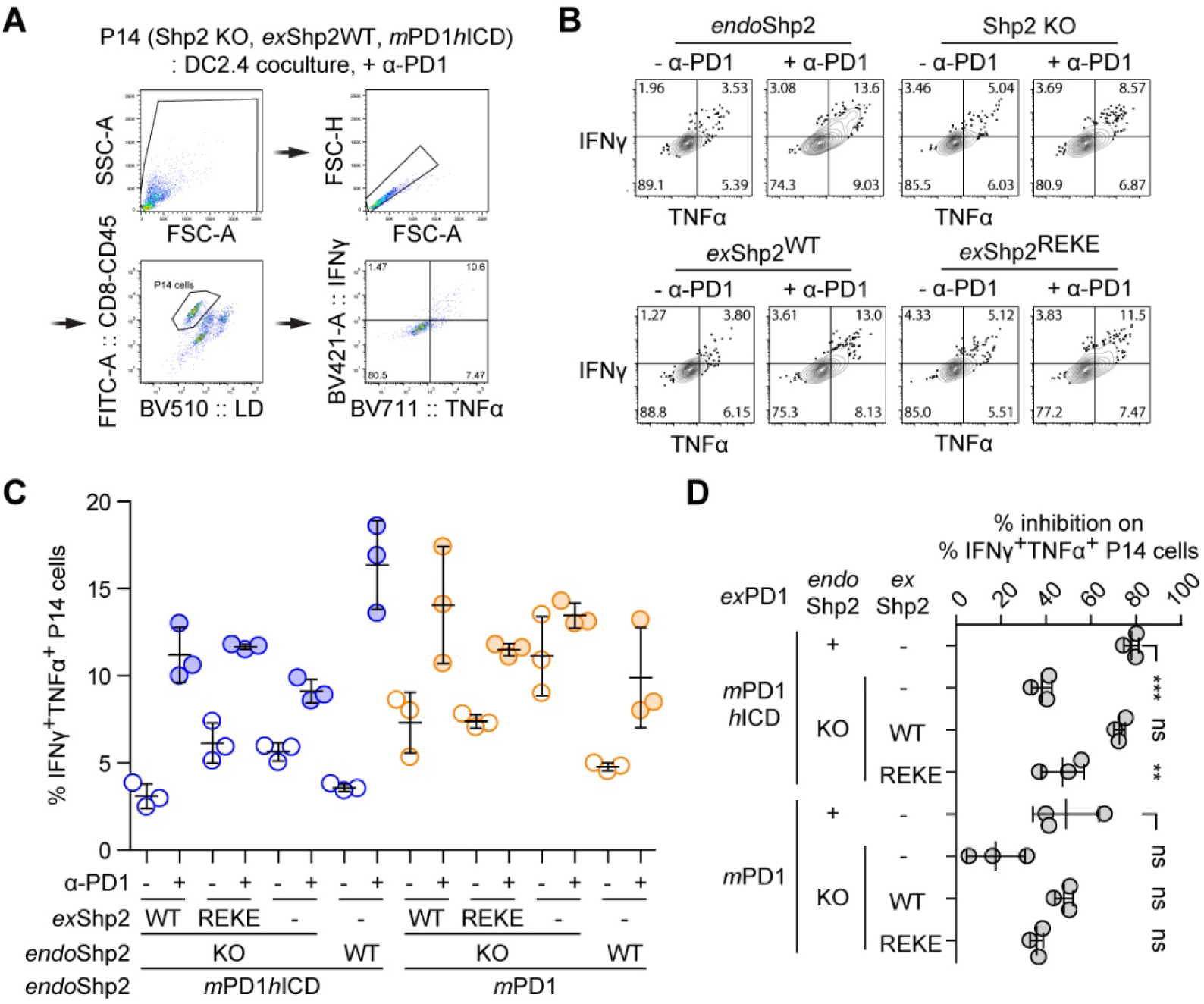
Shp2 REKE mutation attenuates PD1 inhibitory function in mouse primary CD8^+^ T cells. **(A)** Flow cytometry gating strategies to measure %IFNγ^+^TNFα^+^P14 cells in P14: DC2.4 cocultures. **(B)** Representative flow cytometry plots showing the IFNγ and TNFα intracellular staining in the indicated P14 cells in P14: DC2.4 cocultures. **(C)** Dot plots showing %IFNγ^+^TNFα^+^P14 cells under the indicated conditions calculated from B. **(D)** PD1-mediated % inhibition on % IFNγ^+^TNFα^+^ P14 cells under the indicated conditions calculated from C. Data are mean ± SD. **P < 0.01; ***P < 0.001; ns, not significant; student’s t-test.

**Fig. S9.**
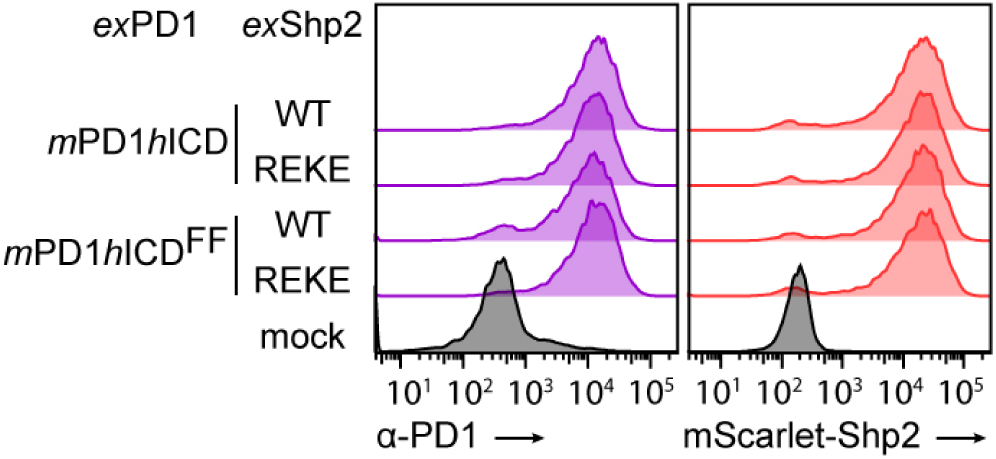
Relevant protein expressions in P14 cells used for the adoptive transfer melanoma model. Flow cytometry histograms showing the expression of *ex*PD1 (left) and *ex*Shp2 (right) in the indicated P14 cells.

## Materials and Methods

### Reagents

DMEM (Dulbeccos Modification of Eagles Medium, [+] 4.5 g/L glucose, L-glutamine [-] sodium pyruvate, #10-017-CM) and RPMI-1640 (Roswell Park Memorial Institute, [+] L-glutamine, 25 mM HEPES, #10-041-CM) media were purchased from Corning. FBS (Fetal Bovie Serum, #FB-02) was purchased from Cell Culture Collective, Inc. 100x P/S (Penicillin-Streptomycin solution, #SV30010) was purchased from Cytiva. Polyethyleneimine (PEI-25000, #NC1014320), 1x PBS (Phosphate Buffered Saline, #10010049), Saponin (#558255100GM), recombinant mouse IFNγ (#315-05) and Foxp3 / Transcription Factor Staining kit (#00-5523-00) were purchased from Fisher Scientific. PFA (Paraformaldehyde, #15714) was purchased from Electron Microscopy Sciences. SHP099 (#S8278) and TNO155 (#S8987) were purchased from Selleck Chemicals.

Mojosort mouse CD8^+^ naïve T cell isolation kit, Human IL-2 ELISA kit (#431816) and BFA (Brefeldin A, #420601) were purchased from Biolegend. SEE (Staphylococcal Enterotoxin E, #ET404) was purchased from Toxin Technologies. gp_33-41_ peptide (Lymphocytic Choreomeningitis Virus Glycoprotein 33-41, #AS-61669) was purchased from AnaSpec. Recombinant human IL-2 (#200-02) was purchased from PeproTech. Snap-Cell505 star (SC505: #S9103S) was purchased from NEB. Janelia Fluor 646 maleimide (JF646-maleimide) was purchased from Bio-Techne (TOCRIS). SNAP-ligand Janelia Fluor 549 was provided by Janelia Research Campus. Hepes (#H3375), hellmanex (#Z805939), D-(+)-glucose (#G8270), imidazole (#I202), sodium chloride (#746398), sodium hydroxide (#S5881), sodium phosphate dibasic dihydrate (#71643), magnesium chloride (#M2670), potassium chloride (#746436), Duolink® flowPLA Detection Kit – FarRed (DUO94004-40TST), Duolink® In Situ PLA® Probe Anti-Mouse MINUS (DUO92004-30RXN), Duolink® In Situ PLA® Probe Anti-Rabbit PLUS (DUO92002-30RXN) were purchased from Sigma-Aldrich. Calcium chloride was purchased from VWR (CaCl_2_, #97061-904). Ni-NTA resin (#88223), β-mercaptoethanol (#21985023), MEM nonessential amino acids solution (NEAA: #11140050), and sodium pyruvate (#11360070) were purchased from Thermo Fisher Scientific. Glutathione Agarose resin (#G-250-10) was purchased from GoldBio. Polybrene (#TR-1003-G) was purchased from Thomas Scientific.

### Mice

C57BL/6 mice were bred in house. Cas9^+^ C57BL/6 mice were procured from the Jackson Laboratory (B6(C)-Gt(ROSA)26Sorem1.1(CAG-cas9*,-EGFP)Rsky/J), and crossed with P14^+^ transgenic mice to generate Cas9^+/−^ P14^+^ mice, or crossed with *Pdcd1*^−/−^ P14^+^ mice (provided by Dr. Rafi Ahmed at Emory University) to generate *Pdcd1*^−/−^ Cas9^+/−^ P14^+^ mice in house. All animals were housed under specific pathogen–free conditions at the University of California San Diego (UCSD). All procedures were previously reviewed and approved by the UCSD Institutional Animal Care and Use Committee under protocol S18078.

### Mouse primary T cells

Spleens were harvested from Cas9^+/−^ P14^+^ or *Pdcd1*^−/−^ Cas9^+/−^ P14^+^ mice and mashed on 70 μm cell strainer in 1x PBS to collect single cell suspensions of splenocytes. Naïve CD8^+^ T cells were isolated from the splenocytes using MojoSort mouse CD8^+^ naïve T cell isolation kit (BioLegend), mixed with 1 μg/mL α-CD3ε/CD28 in primary cell culture media (RPMI-1640 supplemented with 5% FBS, 1 mM sodium pyruvate, 1x Penicillin-Streptomycin, 1x non-essential amino acid solution, and 55 μM β-mercaptoethanol), and primed in a goat IgG fraction-α-mouse IgG-coated plate at 37°C/5% CO_2_ for 24 h.

### Cell lines and cultures

Jurkat E6-1 cells (#TIB-512) were purchased from ATCC. HEK293T and Raji B cell lines were originally obtained from the laboratory of Dr. Ronald D. Vale at the UC San Francisco, and are maintained in the Hui laboratory at UC San Diego. DC2.4 cells (*51*) were purchased from Millipore-Sigma. B16-gp33 cell line was provided from the laboratory of Dr. Stephen Hedrick at the University of California San Diego. HEK293T cells were maintained in DMEM supplemented with 10% FBS and 1x P/S. Jurkat cells and Raji B cells were maintained in RPMI-1640 media supplemented with 10% FBS and 1x Penicillin-Streptomycin. DC2.4 cells and B16-gp33 cells were maintained in RPMI-1640 supplemented with 10% FBS, 1 mM sodium pyruvate, 1x P/S, 1x non-essential amino acid solution, and 55 μM β-mercaptoethanol. All cells were cultured in a 37°C/5% CO_2_ incubator.

### Gene transduction and knockout in primary cells and cell lines

Jurkat cells and Raji B cells were lentivirally transduced with the gene of interest (GOI) as previously described (*42*). Briefly, GOI was cloned into a pHR vector and co-transfected into HEK293T cells together with lentivirus packaging plasmids (pMD2.G and psPAX2) using PEI. After 72 h, HEK293T cell supernatants were collected, mixed with the target cell lines, and centrifuged at 800 x g for 60 min at 32°C. The target cell lines were cultured in the growth media for 3-5 days, after which transgene expression was confirmed by measuring the fluorescence or antibody stain on flow cytometry. Gene knockout in cell lines was performed as previously described (*42*). Briefly, the target cell lines were electroporated with pX330 vectors encoding both sgRNA and Cas9-GFP and cultured for 2-3 days. PD1KO Jurkat cells were stained with α-PD1, and sorted for the GFP-positive, α-PD1-negative population. Shp2KO Jurkat cell line was cloned via single-cell sorting and validated for Shp2KO via western blotting.

Mouse primary CD8^+^ T cells were transduced with retrovirus to express or knockout PD1 and Shp2, as previously described (*42*). Briefly, pQCX plasmids encoding (1) U6 promoter-SV40-exogenous PD1-2A-Thy1.1 or (2) SV40-exogenous Shp2-mScarlet-I were co-transfected with pCL-Eco packaging plasmid into HEK293T cells to produce retrovirus. The retrovirus carrying the indicated genes were mixed with the primed mouse CD8^+^ T cells in the presence of 10 μg/mL polybrene and centrifuged at 800 x g for 90 min at 32°C. After centrifugation, the medium was replaced with the primary cell culture medium supplemented with 100 U/mL human IL-2, and the cells were cultured at 37°C/5% CO_2_. Transduction and/or knockout efficiencies of PD1 and Shp2 were evaluated on day 6 post-transduction via flow cytometry or α-Shp2 western blotting, respectively.

### Antibodies

Biotin α-human CD3ε (clone Okt3, #317320), Alexa Fluor 647 α-human PD1 (clone NAT105, #367419), Alexa Fluor 488 α-mouse CD45 (clone 30-F11, #103121), BV421 α-mouse IFNγ (clone XMG1.2, #505829), and BV711 α-mouse TNFα (clone MP6-XT22, #506349) were purchased from BioLegend. α-mouse CD3ε (clone 145-2C11, #BE00011), α-mouse CD28 (clone 37.51, #BE0015-1), α-human PDL1 (Atezolizumab, #SIM0009), α-mouse PD1 (clone RMP1-14, #BE0146), and α-Trinitrophenol isotype control (#BE0089) were purchased from Bio X Cell. Goat IgG fraction α-mouse IgG (#08670281) was purchased from MPbio. FITC α-mouse CD8a (clone CT-CD8a, #MA5-17597), α-phospho-Tyrosine (clone pY20, #14-5001-82) and α-GFP (#A-6455) were purchased from Thermo Fisher Scientific. α-mouse Shp2 (clone 79, #BDB610621) was purchased from BD. α-beta-Actin (#4967) and α-ALFA-tag (#54963) were purchased from Cell Signaling Technology.

### Recombinant proteins

Streptavidin (#S888) was purchased from Invitrogen. BSA (Bovine Serum Albumin, #A-420-1) was purchased from Goldbio. Recombinant proteins *h*PDL1-His (#10084-H08H), *h*ICAM-1 (#10346-H08H), *h*CD86-His (#10699-H08H), and *h*PVR-His (#10109-H08H) were purchased from Sino Biological. To obtain Shp2 recombinant proteins, full-length human Shp2 (aa 1–593) variant (wild-type, C459S, or R362E/K364E) or mouse Shp2 (aa 1–593) variant (WT or R362E/K364E) was cloned into pET28a vector with an N-terminus 10x histidine-tag followed by a preScission protease recognition site and were expressed in BL21(DE3) strain of *Escherichia coli*. For generating fluorescently labeled Shp2, SNAPf sequence was inserted between the preScission protease recognition site and Shp2. The expressed Shp2 recombinant proteins in bacterial lysates were captured by Ni-NTA agarose, washed with wash buffer (50 mM HEPES-NaOH, pH 8.0, 150 mM NaCl, 30 mM Imidazole, 7 mM β-mercaptoethanol), and eluted with 20 U/mL preScission protease. The eluted Shp2 recombinant proteins were subjected to gel filtration chromatography using a Superdex 200 Increase column (GE Healthcare) in storage buffer (50 mM HEPES-NaOH, pH 7.5, 150 mM NaCl, 1 mM TCEP, 10% glycerol), and the monomeric fractions were harvested. The ICD of human PD1 (*h*PD1^ICD^, aa 194–288) or of mouse PD1 (*m*PD1^ICD^, aa 196–288) was cloned into pGEX-6P2 vector encoding an N-terminal GST-tag followed by a preScission protease cleavage site and an 8x histidine-tag.

Amino acid sequences of wild-type GFP (GFP^WT^(-7)), neutral GFP (GFP(0)), and positively charged GFP (GFP(+7)) were derived from previous study (*52*), and cloned into pGEX-6P2 vector encoding an N-terminal GST-tag followed by a preScission protease cleavage site. The ICD of human CD3ζ (aa 52–164), CD28 (aa 180–220), and TIGIT (aa 163–244) were fused with a C-terminal GFP(0) and cloned into pGEX-6P2 vector encoding an N-terminal GST-tag followed by a preScission protease cleavage site. GST-tagged recombinant proteins were expressed using BL21(DE3) strain, and captured by Glutathione Agarose 4B beads. The beads were washed with wash buffer (50 mM HEPES-NaOH, pH 8.0, 150 mM NaCl, 7 mM β-mercaptoethanol) and incubated with preScission protease for elution. The eluted recombinant proteins were subjected to gel filtration chromatography using a Superdex 75 Increase column (GE Healthcare) in storage buffer (50 mM HEPES-NaOH, pH 7.5, 150 mM NaCl, 1 mM TCEP, 10% glycerol) and monomeric protein fractions were harvested. The Okt3 scFv sequence (from Patent AU2014292924A1) was inserted into a modified pET28a vector containing an N-terminal PelB signal peptide sequence (KYLLPTAAAGLLLLAAQPAMA) and C-terminal 10x histidine-tag, and expressed in BL21(DE3). The expressed Okt3 scFv was extracted via Tris/EDTA/Sucrose periplasmic extraction method (*53*), captured by the Ni-NTA agarose, washed with wash buffer (50 mM HEPES-NaOH, pH 8.0, 150 mM NaCl, 30 mM Imidazole) four times, and eluted with elution buffer ((50 mM HEPES-NaOH, pH 8.0, 150 mM NaCl, 500 mM Imidazole). The eluted Okt3 scFv was subjected to a Superdex 75 Increase column (GE Healthcare) in storage buffer (50 mM HEPES-NaOH, pH 7.5, 150 mM NaCl, 10% glycerol), and the monomeric fractions were harvested. Purified SNAP-tagged Shp2 proteins were incubated with 10-fold excess SC505 or SNAP-ligand Janelia Fluor 549 in the storage buffer for 1 h at 23 °C and subjected to a Superdex 200 Increase column to harvest SC505-labeled, monomeric proteins. For preparing pre-phosphorylated ICD proteins, purified ICD proteins were phosphorylated by mixing with 50 nM Lck and 4 mM ATP in phosphorylation buffer (50 mM HEPES-NaOH, pH 7.5, 150 mM NaCl, 1 mg/mL BSA, 10 mM MgCl_2_, 10 mM Na_2_VO_4_) for 3 h at 23 °C, then subjected to Superdex 75 Increase column to collect monomeric fractions. Intact or pre-phosphorylated PD1^ICD^ was fluorescently-labeled by mixing with 1.25-fold excess JF646-maleimide at 23 °C in the storage buffer for 1 h, followed by Superdex 75 Increase column purification to collect monomeric fractions. All purified proteins were snap-frozen, and stored at -80 °C until use.

### Flow cytometry

To analyze protein expression in cell lines or primary T cells, cells were incubated with antibodies diluted in 1x PBS containing 2% FBS for 30 min on ice. The cells were washed and resuspended in 1x PBS containing 2% FBS and analyzed using FACS Fortessa X-20 (BD).

### Lipids

1-palmitoyl-2-oleoyl-glycero-3-phosphocholine (16:0-18:1 PC, POPC, #850457), 1,2-dioleoyl-sn-glycero-3-[(N-(5-amino-1-carboxypentyl)iminodiacetic acid)succinyl] (nickel salt) (18:1 DGS-NTA(Ni), #790404), 1,2-dipalmitoyl-sn-glycero-3-phosphoethanolamine-N-(biotinyl) (sodium salt) (16:0 Biotinyl PE, #870285), 1,2-dioleoyl-sn-glycero-3-phosphoethanolamine-N-[methoxy(polyethylene glycol)-5000] (ammonium salt) (18:1 PEG5000 PE, #880230) were purchased from Avanti Research. SUVs (Small Unilamellar Vesicles) were prepared by freeze-thaw methods (*17*).

### Cell-SLB assay

SLBs were prepared as previously described, with modifications (*11*). Briefly, a glass-bottom 96-well plate (Cellvis) was cleaned with 2.5% Hellmanex for 12 h at 50 °C and extensively washed with ddH_2_O and stored at 23 °C. On the day of experiment, cleaned wells were etched with 6 N NaOH at 50 °C for 2 h, washed with ddH_2_O and 1x PBS, and incubated with SUVs (95.9% POPC, 2% Biotinyl PE, 2% DGS-NTA(Ni), 0.1% PEG5000 PE) at 50 °C for 1 h to form SLBs on glass. SLBs were washed with 1x PBS containing 1 mg/mL BSA thrice, and then incubated with 1 mg/mL streptavidin, 5 nM *h*ICAM-1-His, and 1 nM *h*PDL1-His at RT for 1 h. The SLBs were washed with 1x PBS containing 1 mg/mL BSA to remove unbound proteins, and incubated with 2 µg/mL biotin Okt3 at 23 °C for 30 min. The functionalized SLBs were washed with 1x PBS containing 1 mg/mL BSA and with 1x imaging buffer (20 mM HEPES-NaOH pH 7.5, 137 mM NaCl, 5 mM KCl, 1 mM CaCl_2_, 2 mM MgCl_2_, 0.7 mM Na_2_HPO_4_, 6 mM D-Glucose), then were incubated at 37°C for 15 min before adding cells. Target cells were resuspended in pre-warmed 1x imaging buffer, added to the SLB-containing wells, and incubated at 37°C. Time-lapse imaging and FRAP experiments (details described below) were performed at 37°C. For measuring PD1 microcluster indices and Shp2 recruitment, cells were fixed with 2% PFA at 5 min after adding to SLBs, incubated for 10 min at RT, and washed with 1x PBS containing 1 mg/mL BSA. Cells landing on SLBs were imaged using an Eclipse Ti2-E microscope (Nikon) equipped with a CSU-W1 SoRA spinning disk unit (Yokogawa) and controlled by NIS-Elements software (Nikon), with either a Plan Apo ʎ 60x NA-1.4 oil immersion objective (WD 130 µm, for fixed cells) or a Plan Apo ʎ 100x NA-1.45 oil immersion objective (WD 130 µm, for live cells).

### PLA-SLB assay

SLBs were formed and washed as described above. To present the corresponding ligands, the SLBs were incubated at 23 °C for 1 h with the following His-tagged protein mixtures: (1) 1.5 nM Okt3 scFv-His, 5 nM *h*ICAM-1-His, and 1 nM *h*PDL1-His for PD1:CD3ζ PLA; (2) 1.5 nM Okt3 scFv-His, 5 nM *h*ICAM-1-His, 1 nM *h*PDL1-His, and 1 nM *h*CD86-His for PD1:CD28 PLA; and (3) 1.5 nM Okt3 scFv-His, 5 nM *h*ICAM-1-His, 1 nM *h*PDL1-His, and 1 nM *h*PVR-His for PD1:TIGIT PLA. The functionalized SLBs were washed with 1x PBS containing 1 mg/mL BSA and with 1x imaging buffer, followed by incubation at 37 °C for 15 min. Jurkat cells were resuspended in pre-warmed 1x imaging buffer, added to the SLB-containing wells, and incubated at 37 °C for 5 min. After the incubation, cells were fixed with 2% PFA and permeabilized with 1x PBS containing 20 mg/mL BSA and 0.1% Saponin, then incubated with α-GFP and α-ALFA-tag at 4°C for 12 h. PLA for α-GFP and α-ALFA-tag was performed using the Duolink® flowPLA Detection Kit (FarRed) together with the Duolink® In Situ PLA® Probe Anti-Mouse MINUS and Duolink® In Situ PLA® Probe Anti-Rabbit PLUS (Sigma-aldrich), following the manufacturer’s protocol. Cells were then imaged using an Eclipse Ti2-E microscope (Nikon) equipped with a CSU-W1 SoRA spinning disk unit (Yokogawa) and controlled by NIS-Elements software (Nikon), with a Plan Apo ʎ 60x NA-1.4 oil immersion objective (WD 130 µm).

### In vitro condensation assay

Recombinant Shp2 and/or PD1^ICD^ proteins were mixed at the indicated concentrations with LLPS buffer (50 mM HEPES-NaOH, pH 7.5, 125 mM NaCl, 2.5 mM MgCl_2_, 1 mM TCEP, 5% glycerol, 8% PEG 8000) in Eppendorf tubes and incubated for 5 min at 23 °C. The samples were then transferred to a 384-well glass-bottom plate (Cellvis) and centrifuged at 500 x g for 1 min at 23 °C. Optical density at 600 nm (OD_600_) was measured using a Spark multimode microplate reader (Tecan) both before and after the sample transfer to determine background signal and protein turbidity. Protein condensates landing on the glass surface were imaged using an Eclipse Ti2-E microscope (Nikon) equipped with a CSU-W1 SoRA spinning disk unit (Yokogawa) and, a Plan Apo ʎ 100x NA-1.45 oil immersion objective (WD 130 µm), controlled by NIS-Elements software (Nikon). To examine the effect of salt on Shp2 condensation, Shp2 and PD1^ICD^ were mixed into LLPS buffer containing varying concentrations of NaCl (**Fig. 5B,C**). Alternatively, Shp2:PD1ICD condensates were first formed in LLPS buffer with 125 mM NaCl, then mixed with 125 mM or 5 M NaCl to achieve the indicated final NaCl concentrations (**Fig. 5D**). To examine receptor ICD enrichment, 8 μM Shp2, 8 μM PD1^ICD^, and 50 nM receptor ICD or GFP(0) were mixed into LLPS buffer, incubated at 23 °C for 5 min before transferring into PF127-treated glass-bottom wells (*54*).

### FRAP assays

The fluorescent images were acquired at a minimum of three timepoints before photobleaching to establish a baseline. The region of interest (ROI) was photobleached using a 405 nm laser until fluorescence within the ROI decreased by more than 70%. Post-bleaching images were acquired to capture the recovery phase. Image analysis was conducted using Fiji (*55*). To obtain the fluorescent time-course, fluorescent intensity of the photobleached protein condensate was measured at each time point and normalized to that of a neighboring, unbleached condensate. The resulting fluorescent recovery curves were fit to a “One-phase association” model using GraphPad Prism Version 5.01 (GraphPad Software, Inc.) to calculate the recovery time constant τ.

### Microcluster image analysis

Image analysis was conducted using Fiji (*55*). Image areas containing single cell were cropped from the raw images. Cropped raw cell images were processed to create mask images for either whole cell area or microcluster area. To generate mask images for whole cell area, cropped images were background-subtracted using the “MinError” threshold filter. To create mask images for microclusters, cropped raw cell images were processed in two background subtraction steps. First, a background image was generated by applying the median filter (sigma=10) and subtracted from the original image to remove the broad background signals. Second, residual background was further removed using the “Otsu” threshold filter. Cropped raw cell images were applied to the corresponding whole cell area mask and microcluster mask. The mean fluorescence of the whole cell area was measured to determine the synaptic fluorescent intensities of PD1 or Shp2. The microcluster indices for each cropped cell image was calculated by dividing the fluorescent intensity of a microcluster image by the one of a whole cell area image.

### PTPase activity assay

To measure the Shp2 PTPase activity using DiFMUP in **Fig. 6 and Fig. S2**, 20 nM of *h*Shp2^WT^, *h*Shp2^REKE^, or *h*Shp2^C459E^ was mixed with the indicated concentration of p-*h*PD1^ICD^ in HEPES buffered saline (50 mM HEPES, pH 7.5, 150 mM NaCl, 1 mM TCEP) and incubated at 23 °C for 1 min, followed by the addition of DiFMUP at the indicated concentrations. The FI of DiFMU, a dephosphorylated product of DiFMUP, was continuously monitored for 60 min at 23 °C using a Spark multimode microplate reader (Tecan). DiFMU concentrations at each time point were determined from a standard curve generated by plotting the FI plateau against the initial DiFMUP concentrations. The initial reaction velocity (V0) was calculated as the slope of a linear fit of the first 30 s of data points divided by the Shp2 concentration. The V0-[DiFMUP] plot was fitted using the “Michaelis-Menten” model using GraphPad Prism 5.0. To assess the Shp2 PTPase activity toward p-*h*CD3ζ, the indicated concentrations of Shp2 variants and p-*h*PD1^ICD^ were incubated with 0.5 μM p-*h*CD3ζ in LLPS buffer at 23 °C for the indicated durations. Reactions were terminated by mixing with 33% volume of 4x SDS sample buffer and subjected to α-pY IB, followed by chemiluminescent imaging using a ChemiDoc system (Bio-Rad). The optical density of p-*h*CD3ζ bands was quantified using Fiji (*55*) and normalized to that at time zero. The time course of % p-*h*CD3ζ was fitted to the “One phase decay” model using GraphPad Prism 5.0.

### Jurkat:Raji coculture assay

To evaluate PD1 inhibitory function in Jurkat cells in **Fig. S4**, Jurkat cells expressing the indicated PD1 and Shp2 were mixed with SEE-loaded, CD80 KO Raji cells expressing human PDL1-mCherry in the absence or presence of α-human PDL1 blockade, and incubated at 37°C/5% CO_2_ for 6 h. The IL-2 secreted in the supernatant was quantified using a human IL-2 ELISA kit. The use of CD80-deficient Raji cells prevented cis-interactions between CD80 and PDL1 (*56–58*), which could otherwise mask PD1–PD-L1 engagement.

### P14:DC2.4 coculture assay

To evaluate PD1 inhibitory function in mouse primary T cells, P14 cells transduced with the indicated sgRNA and/or exogenous genes were generated as described above. DC2.4 cells were treated with 30 U/mL mouse IFNγ at 37°C/5% CO_2_ for 20 h to induce *m*PDL1 expression. *m*PDL1-expressing DC2.4 cells were then incubated with the indicated concentration of gp_33-41_ peptide at 37°C/5% CO_2_ for 1 h, and washed with pre-warmed 1x PBS.

P14 cells were incubated with α-mouse PD1 blockade or isotype control at 37°C/5% CO_2_ for 30 min, added to PDL1-expressing, gp_33-41_-pulsed DC2.4 cells, and cultured at 37°C/5% CO_2_ for 4.5 h in the absence or presence of BFA for analyzing cytokine secretion or intracellular cytokine production, respectively. The concentration of secreted IFNγ was measured using ELISA MAX™ Standard Set Mouse IFN-γ (BioLegend). For quantifying intracellular cytokine production, cells were stained with Zombie Aqua Fixable Viability Dye, and then with Alexa Fluor 488 α-mouse CD8a and FITC α-mouse CD45. The stained cells were fixed and permeabilized using Foxp3 / Transcription Factor Staining kit, and further stained with BV421 α-mouse IFNγ and BV711 α-mouse TNFα. The stained cells were fixed with 2% PFA, and analyzed using FACS Fortessa X-20 (BD).

### Adoptive T cell transfer melanoma model

To assess PD1 inhibitory function in mouse primary T cells *in vivo*, P14 cells were isolated from female *Pdcd1*^−/−^ Cas9^+^ P14 C57BL/6 mice, transduced with the indicated gRNA and exogenous genes generated as described above, and expanded *in vitro* for seven days prior to adoptive transfer.

Male C57BL/6 mice (7-10 weeks old) were subcutaneously inoculated with 2.5 × 10^5^ B16-gp33 melanoma cells in the right flank. Seven days after tumor inoculation, 2.5 × 10^5^ P14 cells were transferred intravenously via retro-orbital injection under isoflurane anesthesia. Tumor growth was monitored three times per week, and tumor volume calculated using the formula: (length x width^2^) / 2. Mice were randomized based on the cage and tumor size on the day of T cell transfer.

